# Genetic immune escape landscape in primary and metastatic cancer

**DOI:** 10.1101/2022.02.23.481444

**Authors:** Francisco Martínez-Jiménez, Peter Priestley, Charles Shale, Jonathan Baber, Erik Rozemuller, Edwin Cuppen

**Author notes:** Corresponding authors, Francisco Martínez-Jiménez Edwin Cuppen. These authors contributed equally to this work (Francisco Martínez-Jiménez, Peter Priestley).

## Abstract

Immune surveillance escape is a hallmark of tumorigenesis^1^. Multiple studies have characterized the immune escape landscape across several untreated early-stage primary cancer types^2–4^. However, whether late-stage treated metastatic tumors present differences in genetic immune escape (GIE) prevalence and dynamics remains unclear. Here, we performed a pan-cancer characterization of GIE prevalence across six immune escape pathways in 6,457 uniformly processed Whole Genome Sequencing (WGS) tumor samples including 58 cancer types from 1,943 primary untreated patients and 4,514 metastatic patients. To effectively address the complexity of the Human Leukocyte Antigen (HLA-I) locus and to characterize its tumor status, we developed LILAC, an open-source integrative framework. We demonstrate that one in four tumors harbor GIE alterations, with high mechanistic and frequency variability across cancer types. GIE prevalence is highly consistent between primary and metastatic tumors for most cancer types with few exceptions such as prostate and thyroid carcinomas that have increased immune evasion frequencies in metastatic tumors. Positive selection analysis revealed that GIE alterations are frequently selected for in tumor evolution and that focal LOH of HLA-I, unlike non-focal LOH of HLA-I, tends to lose the HLA allele that presents the largest neoepitope repertoire. We also unraveled tumor genomic features contributing to immune escape incidence, including DNA repair deficiency, APOBEC activity, tobacco associated mutation load and viral DNA integration. Finally, there is a strong tendency for mid and high tumor mutation burden (TMB) tumors to preferentially select LOH of HLA-I for GIE whereas hypermutated samples favor global immune evasion strategies. Our results indicate that genetic immune escape is generally a pre-metastatic event during tumor evolution and that tumors adapt different strategies depending on their neoepitope burden.

## Introduction

Cancer immune escape is the process whereby tumor cells prevent their elimination by the immune system^5^. This process is leveraged by tumor cells as a response to the accumulation of tumor specific alterations, which are susceptible to be presented -in the form of neoepitopes-by the Major Histocompatibility Complex-I (MHC-I). Despite the significant advances in predicting which neoantigens are presented as neoepitopes, it is still challenging to accurately select the neoepitopes capacity to trigger cytotoxic T-cell responses (i.e., the neoantigens) based on genomics data^6^.

Tumorigenesis often selects genomic somatic alterations in order to hinder the recognition and/or elimination by the immune system^7^, a process hereafter referred to as genetic immune escape (GIE). Such alterations operate through different mechanisms, including partial or complete abrogation of neoepitope presentation^8^ or suppression of pro-apoptotic signals from the surrounding immune cells^9^. Therefore, identification of GIE events across human cancers is key to understanding the interplay between cancer cells and the immune system as well as to enable effective precision medicine based on immunotherapy.

Prior studies have performed cancer type specific molecular profiling of GIE events and their phenotypic implications in certain cancer types, including seminal work in non-small cell lung cancer^2, 3^ (NSCLC) and colorectal carcinoma^4^, among others^10, 11^. Others have performed an extensive analysis of LOH HLA-I across thousands of tumor samples^12^. However, a pan-cancer analysis of the prevalence and impact of diverse GIE events is currently lacking. In addition to that, the focus of these studies was to portray GIE in early-stage untreated primary tumors, whereas the changes induced by exposure to treatment and by the metastatic bottleneck have not been comprehensively addressed but might be highly relevant for advanced cancer treatment.

One of the main challenges to perform such analyses lies in the extraordinary diversity of the HLA locus, consisting in multiple genes and different maternal and paternal make ups. To date, more than 15,000 different sequences of the HLA-A, HLA-B and HLA-C genes have been reported^13^, which significantly hampers the identification of tumor specific somatic alterations. This prompted the development of tools that specifically identify LOH of HLA class I alleles (LOH HLA-I)^14^ or somatic mutations^15^ from whole exome/genome sequencing data. These studies revealed the relevance of assessing tumor specific HLA alterations in tumorigenesis. However, none of these tools provide an integrative characterization of the HLA-I tumor status, which includes HLA-I typing, allelic imbalance, LOH-HLA-I and somatic mutation annotation.

In this study we present a pan-cancer landscape of the GIE prevalence in primary (represented by the PCAWG cohort) and metastatic patients (represented by the Hartwig cohort). Furthermore, to address the complexity of the HLA-I locus, we developed LILAC, an open-source integrative framework that characterizes the HLA-I locus, including its tumor status from WGS data. We applied LILAC and a universal tumor processing pipeline to establish a comprehensive portrait of GIE events and their positive selection landscape across six different pathways recurrently associated with an immune evasion phenotype: aberration of the HLA-I locus itself, mutations in the antigen presentation machinery, IFN-γ signaling inactivation, PD-L1 amplification, CD58 inactivation and epigenetic immune escape (Fig. 1a and Supp. Table 1; methods GIE alterations). We also studied how the TMB and other genomic and environmental features influences the prevalence of GIE alterations, providing insights into tumor evolution and its interplay with the immune system.

**Figure 1.**
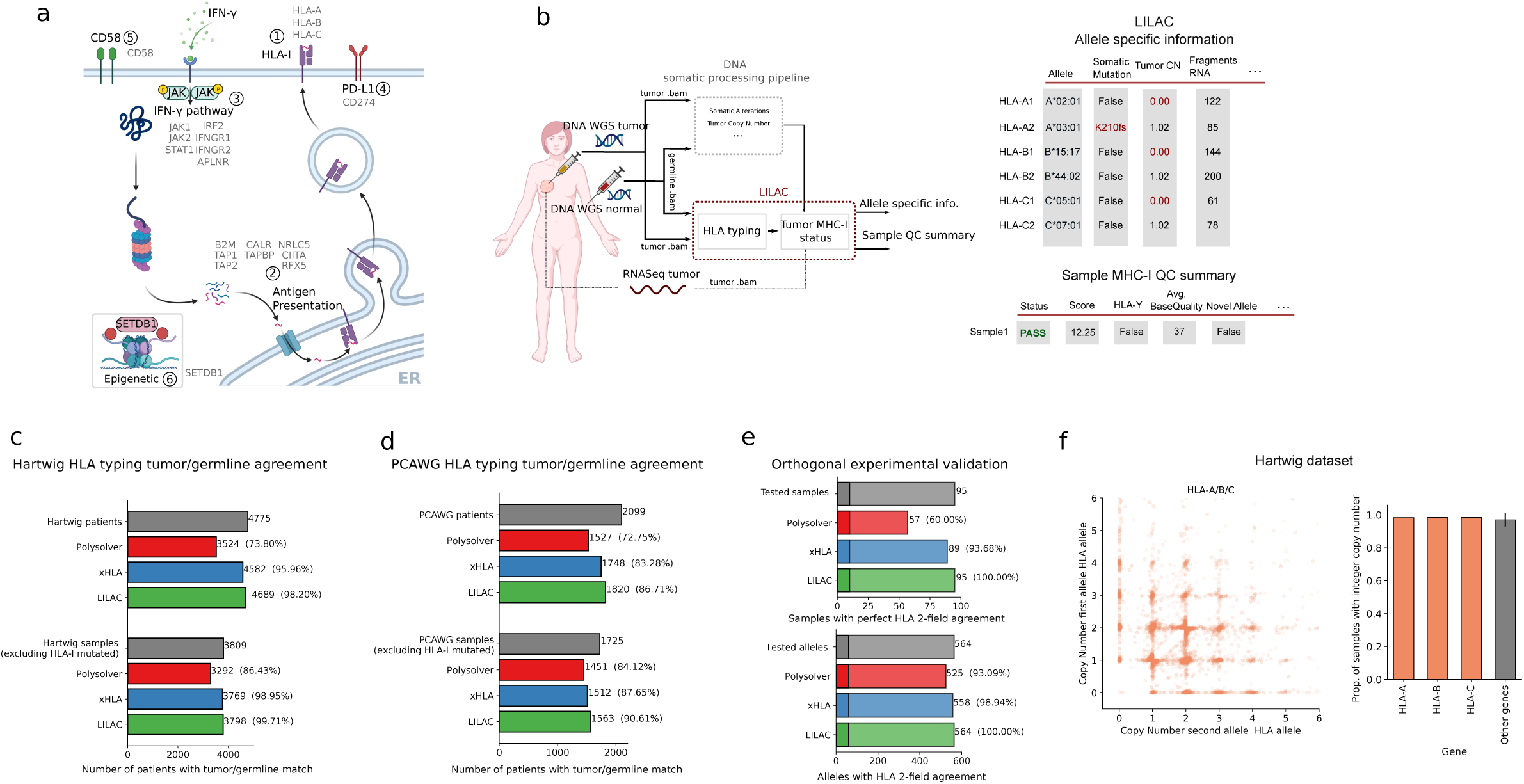
Inference of HLA-I tumor status with LILAC. a) Representation of the six immune escape pathways considered in this study alongside their associated genes. Adapted from “MHC Class and II Pathways”, by BioRender.com (2021). b) Left, workflow of the tumor analytical pipeline including LILAC. Right, tables show an illustrative example of LILAC’s patient-specific reports. Partially created with BioRender.com. c) HLA-I typing tumor and germline agreement in Hartwig and d) PCAWG. e) LILAC’s experimental validation. f) Left, copy number of minor and major alleles of HLA-A, HLA-B and HLA-C in Hartwig. Right, proportion of samples with integer copy number of HLA-I genes (orange) compared to the average of 1,000 random genes in Hartwig (gray). The error bar represents the standard deviation.

## Results

### Inference of HLA-I tumor status with LILAC

Inference of the correct HLA-I tumor status is fundamental to identify GIE alterations (Fig. 1a), to estimate the neoepitope repertoire and burden and to predict the response to Immune Checkpoint Inhibitors^16, 17^ (ICI). None of currently available HLA typing tools provides an integrated characterization of the germline and the tumor HLA-I locus. To bridge this gap, we developed LILAC, a framework that performs HLA-I typing for the germline of each patient as well as determining the status of each of those alleles in the tumor including complete loss of one or more alleles, allele specific somatic mutations and allelic imbalance using whole genome sequencing data of tumor-normal pairs as input. LILAC is also able to detect novel HLA-I alleles. Finally, it provides allele-specific and sample level quality control measurements, which assist in the interpretation of LILAC’s output (Fig. 1b, see methods LILAC and Supplementary Note 1).

We first assessed LILAC’s HLA-I typing robustness by independently calculating the germline and tumor HLA-I 2-field calling agreement across 6,874 patients (see methods LILAC), including 4,775 patients from the Hartwig^18^ dataset and 2,099 from the PCAWG^19^ cohort. LILAC showed the highest agreement (98.20% and 86.71% in Hartwig and PCAWG, respectively) compared to two state-of-the-art HLA typing tools, Polysolver^15^ and xHLA^20^ (Fig. 1c and Fig.1d top panels and Supp. Table 2). The Hartwig dataset showed higher normal-tumor agreement for all tools, possibly due to the higher sequencing coverage and read quality of this dataset which was generated with more recent sequencing platforms. The exclusion of samples bearing HLA-I somatic alterations -that may hinder the comparison-rendered a very high agreement for LILAC across both datasets (99.71% for Hartwig and 90.61% for PCAWG), which was consistently higher than the two other methods on the same set of samples (Fig. 1c and Fig. 1d, bottom panels and and Supp. Table 2). In a three-way comparison, LILAC also displayed the highest overlap with the predictions from the other tools across both datasets (Supp. Fig. 1a and Supp. Fig. 1b). Next, we evaluated LILAC HLA-I typing sensitivity in a set of 95 samples -including 10 from tumor biopsies-with an independent orthogonal and clinically validated HLA-I typing approach by high-to allelic resolution HLA typing (see methods LILAC experimental validation). LILAC showed a perfect 100% 2-field agreement across the 564 alleles, higher than Polysolver (93.09%) and xHLA (98.94%) agreements (Fig. 1e. and Supp. Table 2). Moreover, LILAC reported nine somatic mutations in seven of the tumor biopsies evaluated. All of them (100%) were perfectly matched by the orthogonal approach (Supp. Table 2), highlighting our framework’s ability to identify and map somatic variants into HLA alleles in tumor samples.

HLA allele specific tumor copy number (CN) determination is key to identify LOH of HLA-I genes in tumors, a well-established mechanism of immune evasion^12, 14^, as well as to determine allelic imbalance events. LILAC annotates allele specific ploidy levels of each HLA-I allele based on the purity corrected local tumor copy number estimations and the number of fragments assigned to each allele (see methods LILAC). WGS data provides adequate resolution to annotate purity-adjusted minor and major allele ploidy in the HLA-I locus, illustrated by the similar proportion of integer copy number of HLA-I genes compared to a random set of 1,000 genes (Fig. 1f and Supp. Fig. 1c). Moreover, the three tumor samples harboring LOH of HLA-I according to our framework and evaluated by the orthogonal approach displayed a strong allelic imbalance according to the experimental validation (Supp. Table 2).

In summary, LILAC shows a highly sensitive and specific HLA-I typing performance and an excellent ability to map tumor somatic events into the HLA alleles in WGS samples with sufficient quality and thus forms an excellent basis for neoepitope prediction and studying immune evasion mechanisms.

### Genetic Immune Escape (GIE) in primary and metastatic tumors

In addition to the HLA-I targeting alterations (i.e., LOH of HLA-I, somatic mutations and homozygous deletions), other tumor somatic alterations may lead to an immune evasion phenotype. Therefore, we combined LILAC with the Hartwig analytical comprehensive cancer WGS pipeline^18, 21^ to annotate GIE events across six pathways strongly associated with immune escape (Fig. 1a and Supp. Table 1; methods GIE alterations) across 6,457 uniformly processed WGS samples, including 1,943 untreated primary patients from PCAWG and 4,514 metastatic patients from Hartwig (Fig. 2a, Suppl. Fig. 2a and Suppl. Table 3; methods Data collection and processing). In total, these patients were classified into 58 cancer types, which included 38 tumor types with sufficiently high representativeness (i.e., number of patients greater or equal than 15) in the metastatic cohort, 27 in the primary dataset and 20 cancer types with sufficient representation in both datasets to allow for a comparison between primary and metastatic tumors (Fig. 2b, Supp. Fig. 2b-c and Supp. Table 3).

**Figure 2.**
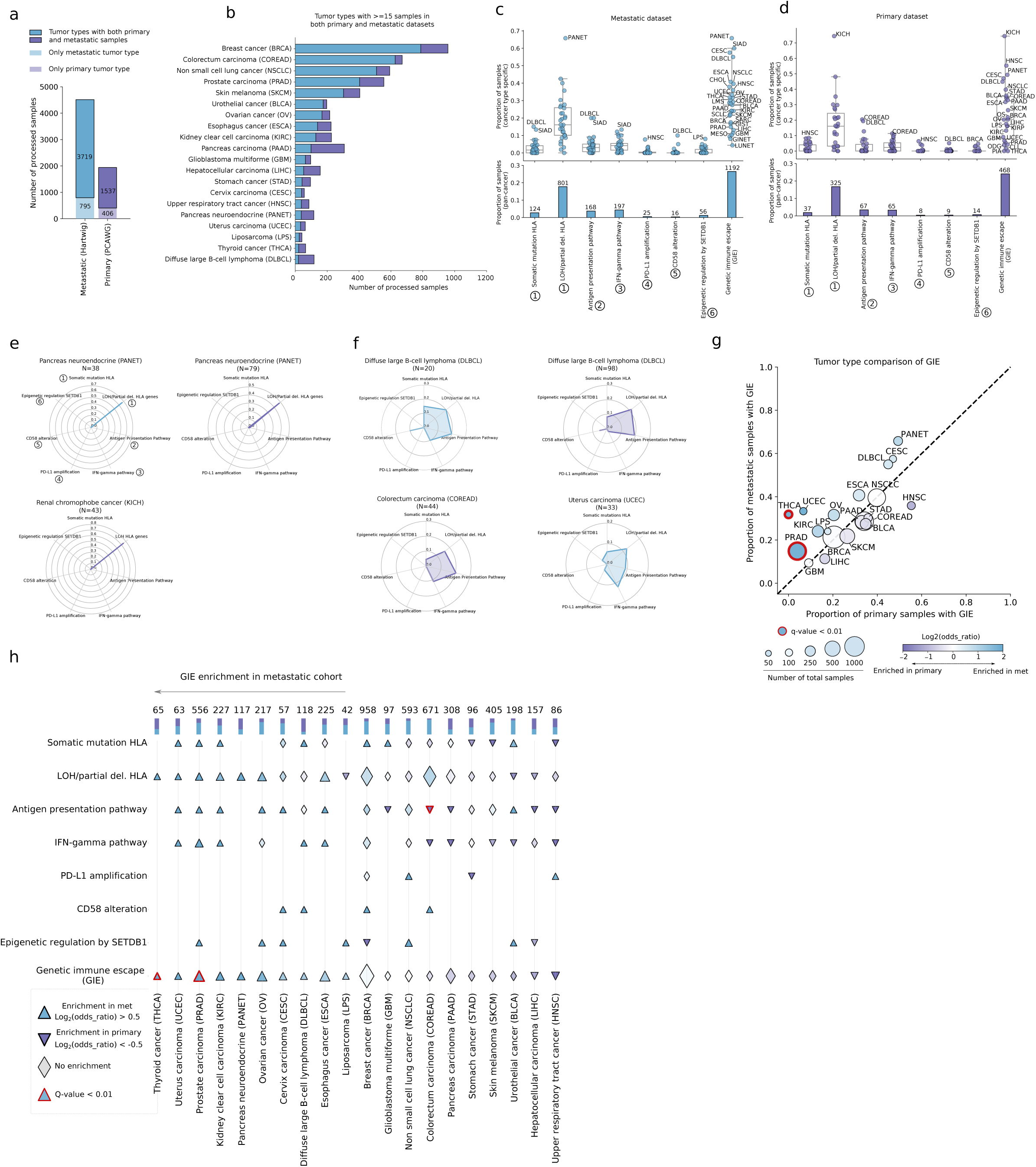
GIE in primary and metastatic tumors. a) Total number of uniformly processed WGS samples included in the study from the metastatic (Hartwig) and primary (PCAWG.) datasets. b) Number of cancer type samples from each dataset across cancer types with at least 15 samples in both datasets. c) top, cancer type specific proportion of metastatic samples with GIE alterations across the six pathways and the combined group. bottom, pan-cancer proportion and number of samples with GIE alterations in the metastatic group. d) analogous for the primary datasets. Box-plots: center line, median; box limits, first and third quartiles; whiskers, lowest/highest data points at first quartile minus/plus 1.5× IQR. The label numbers represent the associated immune escape pathway, relative to Figure 1a. e) and f) Radar plots representing cohort specific proportion of GIE alterations across the six pathways from Fig. 1a. g) Combined proportion of primary (PCAWG) and metastatic (Hartwig) samples affected by GIE alterations across 20 cancer types. Size of dots are proportional to the number of total samples. The color of each dot is proportional to the log_2_(odds ratio). Red edge lines represent an adjusted Fisher’s exact test p-value < 0.01. h) top stacked bars, number and proportion of combined (metastatic and primary) cancer type samples. main panel, pathway specific comparison alongside its significance. SIAD, small intestine cancer. CHOL, cholangiocarcinoma. LMS, leiomyosarcoma. SCLC, small cell lung cancer. SARC, sarcoma various. GIST, gastrointestinal stromal tumor. MESO, mesothelioma. GINET, gastrointestinal neuroendocrine. LUNET, lung neuroendocrine. PIA, Pilocityc astrocytoma. KIRC, Kidney papillary carcinoma. OS, osteosarcoma. ODG, oligodendroglioma. The remaining cancer types acronyms are displayed in panel b).

GIE prevalence showed high mechanistic and frequency variability across primary and metastatic cancer types (Fig. 2c and Fig. 2d top panels; Supp. Table 4). The median proportion of patients harboring GIE alterations per cancer type was 0.27 for the metastatic cohort and 0.20 for primary tumors, both showing highly dispersed distributions (±0.15 std. and ±0.19 std. in metastatic and primary, respectively). In certain cancer types, such as pancreatic neuroendocrine (PANET, metastatic), small intestinal cancer (SIAD, metastatic) and kidney chromophobe cancer (KICH, primary) GIE was present in more than 50% of patient samples (66%, 60% and 74%, respectively) while in others, such as lung neuroendocrine (LUNET, metastatic) and pilocytic astrocytoma (PIA, primary), GIE was extremely rare with 4% and none of the patients, respectively. Overall, one in four patients (26% in metastatic and 24% in primary) presented GIE alterations based on the six investigated pathways (Fig. 2c and Fig. 2d, bottom panels).

The most frequent GIE alteration was partial loss of the HLA-I locus (including both LOH of HLA and homozygous deletions of HLA-I genes and were grouped as LOH of HLA-I for simplicity), which was present in 801 (18%) of metastatic and 325 (17%) of primary cancer patients, followed by IFN-γ inactivation (4% in metastatic and 3% in primary) and antigen presentation pathway (4% in metastatic and 3% in primary). CD58 inactivation was the least frequent immune escape event present in only 16 metastatic and 9 primary patients (Fig. 2c and Fig. 2d, bottom panels). The high GIE rates of KICH and PANET were exclusively due to LOH of HLA-I (Fig. 2e). These LOH events were the result of recurrent chromosome 6 loss^22, 23^ and it is unclear whether LOH HLA-I is the primary driver behind this loss. Other cohorts, such as diffuse large B-cell lymphoma (DLBCL), colorectal carcinoma (COREAD, primary) and uterine carcinoma (UCEC, metastatic) displayed a wider range of GIE mechanisms (Fig. 2f). In summary, GIE prevalence was generally dominated by LOH HLA-I but most cancer types also leverage other mechanisms of immune evasion with lower predominance.

We next sought to investigate whether there was a GIE prevalence difference between primary untreated tumors and late stage metastatic tumors. Comparison by tumor type across the 20 cancer types with sufficient representation (i.e., at least 15 patients in both the primary and metastatic datasets), showed a broad agreement between datasets (Fig. 2g). Although 10 cancer types showed a certain degree of metastatic enrichment (Log_2_(odds ratio) > 0.5; Fig. 2h), only in prostate carcinoma (PRAD) and thyroid cancer (THCA) this difference was statistically significant (Fisher’s exact test corrected p-value < 0.01). Breaking down pathway-specific differences revealed that THCA metastatic enrichment is the result of increased LOH HLA-I incidence, whereas the discrepancies in PRAD are the result of a widespread enrichment across several pathways (Fig. 2h). In general, LOH HLA-I showed a non-significant trend towards metastatic enrichment across 7 of the 10 metastatic enriched cancer types. None of the cancer types showed a significantly higher GIE incidence in primary tumors. Nevertheless, colorectal carcinoma showed more frequent antigen presentation pathway alterations in primary tumors (Fig. 2h), likely associated with higher ratios of hypermutated samples in the PCAWG dataset. In summary, apart from the aforementioned exceptions, the analyzed primary untreated and metastatic cancer types show an overall comparable GIE prevalence, suggesting that immune evasion is generally an early event in tumor evolution that may be further fueled by LOH of HLA-I in later tumorigenic stages.

### Positive selection of HLA-I alterations

We next examined to which extent somatic alterations in HLA-I genes (i.e. HLA-A, HLA-B and HLA-C) were positively selected during tumorigenesis. Since LILAC enables the mapping and consequence type annotation of HLA-I somatic point mutations and indels in HLA alleles, we analyzed their dN/dS ratio^24^ and mutational profile. Pan-cancer grouped HLA-I analysis (see methods Positive selection somatic mutations and indels) exhibited a dN/dS ratio greater than one for nonsense, splice site and truncating variants in both the metastatic and primary datasets (Fig. 3a), showing that these genes are subject to positive selection. Furthermore, pan-cancer and gene-specific dN/dS ratios revealed that HLA-A and HLA-B, but not HLC-C, are positively selected and that they are mostly enriched in truncating variants (frameshifts and nonsense mutations) and not in non-synonymous mutations that change individual amino acids (Fig. 3b and Fig. 3c).

**Figure 3.**
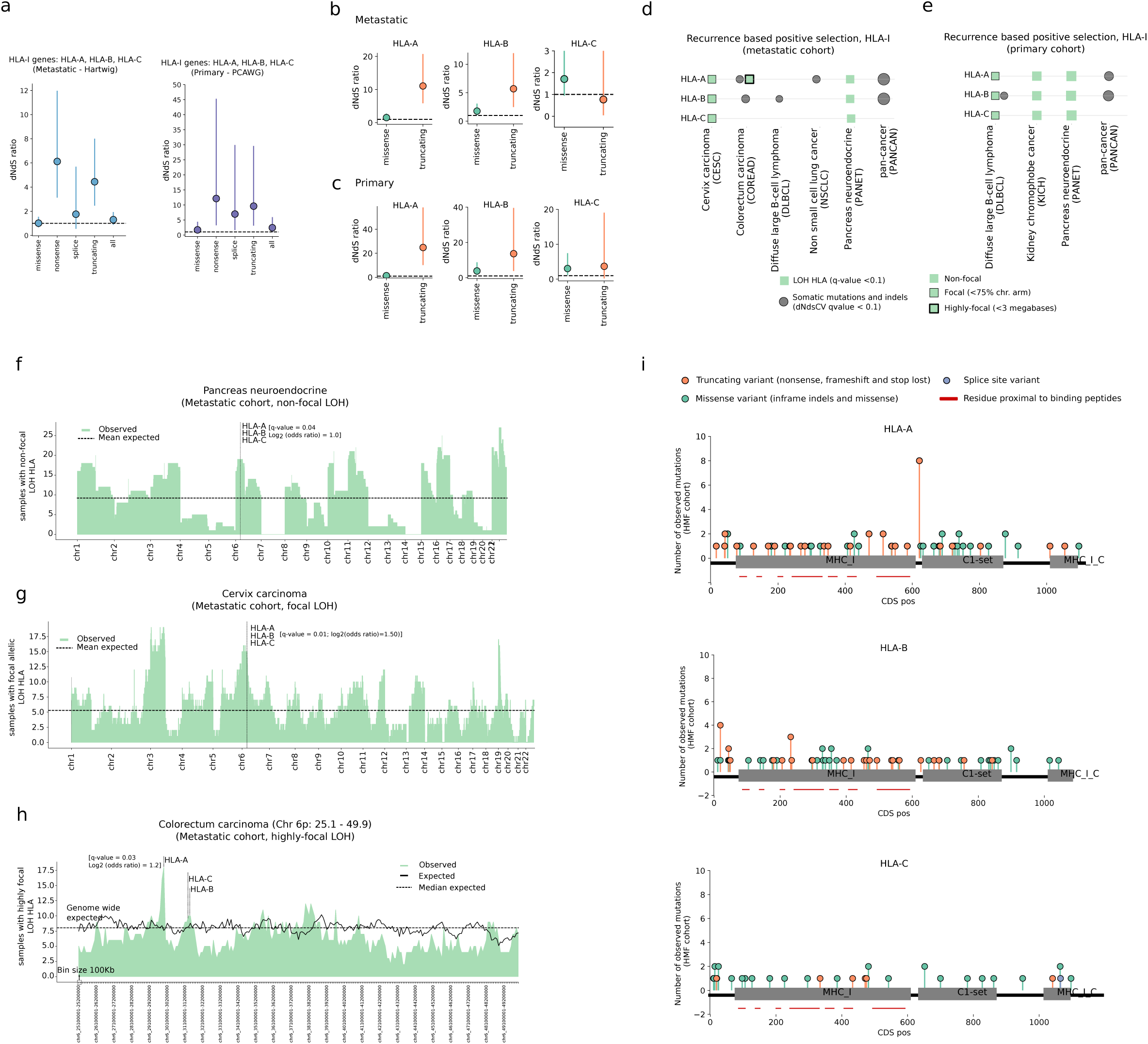
Positive selection of HLA-I. a) Pan-cancer dN/dS ratios of HLA-I genes in the metastatic (left) and primary (right) dataset. Vertical lines represent the 5% and 95% confidence intervals. b) Metastatic pan-cancer and gene-specific dN/dS ratios of HLA-I genes. Vertical lines represent the 5% and 95% confidence intervals. c) Identical for the primary dataset. d) Representation of gene and cancer type specific positive selection of HLA-I in the metastatic and e) primary cohorts. f) Distribution of LOH events along the autosomes in pancreatic neuroendocrine tumors of the metastatic cohort. X-ticks represent the chromosomal starting position. g) Distribution of focal LOH events along the autosomes in cervix cancer tumors of the metastatic cohort. X-ticks represent the chromosomal starting position. h) Distribution of highly-focal LOH surrounding the HLA-I locus spanning from chr6:25.1Mb to chr6:49.9Mb in the metastatic colorectal cancer cohort. Each bin represents 100Kbs. Dashed horizontal lines represent the expected means after randomization. Vertical dashed lines highlight the HLA-I genomic locations. i) Needle plots representing the pan-cancer distribution of somatic mutations along the HLA-A, HLA-B and HLA-C protein sequences in the metastatic dataset. Mutations are coloured according to the consequence type. CDS pos, coding sequence position. Mb, megabase. Kbs, kilobases.

In order to unravel the specificity associated with these global signals we next conducted a gene and cancer type specific analysis of positive selection. We identified positively selected genes from somatic point mutations and indels by using dNdScv and we developed a randomization strategy to assess, at different genomic scales, positive selection in LOH HLA-I, homozygous deletions and copy number amplifications (methods Positive selection and Supp. Table 5). As expected, HLA-A and HLA-B, but not of HLA-C, were deemed as drivers by dNdScv (q-value <0.1) across several cancer types including metastatic colorectal and non-small cell lung cancer, diffuse large B-cell lymphoma as well as the pan-cancer group in both datasets (Fig. 3e and 3f). Since gene inactivation may also be reached by homozygous deletions, we queried the observed versus expected frequency of biallelic HLA-I deletions. We did not observe any biallelic deletion of the entire HLA-I locus (Supp. Table 4). Moreover, none of the gene and cancer-type specific billalelic deletions reached the significance threshold (Supp. Table 5), suggesting that homozygous deletions within the HLA-I might be constrained by purifying selection.

LOH of HLA-I, which is a recurrent genomic event and that had been previously reported as a driver event in non-small cell lung cancer^14^, was more frequently observed than expected across multiple cancer types in both the metastatic and the primary datasets (G-test goodness of fit q-value < 0.1; Fig. 3e and Fig. 3f). More specifically, certain cohorts such as pancreatic neuroendocrine (Fig. 3f) or kidney chromophobe (Supp. Fig. 3a) showed non-focal LOH HLA-I enrichment, which is also compatible with other selective pressures operating at chromosome 6 scale. Others, such as metastatic cervix carcinoma (Fig. 3g) or primary diffuse large B-cell lymphoma, showed focal or highly focal (metastatic colorectal cancer; Fig. 3h) LOH HLA-I patterns, indicative of HLA allele/s loss driven selection.

Finally, somatic point mutations and small indels of HLA-I genes were evenly distributed along their sequences (Fig. 3i and Supp. Fig. 3b). The main exception was the recurrent HLA-A Lys210 frameshift mutation, which was found in 8 MSI metastatic tumors and that overlapped with a homo polymer repeat of CCCCCCC. No enrichment for mutations in amino-acids involved in the peptide binding was observed. Such uniform distribution was in agreement with previous observations^25^ and with the expected profile in tumor suppressor genes dominated by inactivating variants^26^. Furthermore, 33 patients with non-synonymous mutations of HLA-I genes (20% of the total 161 patients with mutations in HLA-I genes) displayed the concurrent loss of the alternative allele by LOH, potentially leading to complete inactivation of the HLA gene. Taken together, our results show that HLA-A and HLA-B somatic inactivation as well as LOH of HLA-I are frequently selected events across several cancer types in tumor evolution whereas complete biallelic deletion of HLA-I and single-amino acid change mutations are rarely exploited by tumors as a mechanism of immune evasion.

### Focal LOH of HLA-I preferentially targets the allele that presents the highest neoepitope repertoire

In HLA-I heterozygous cells, LOH of HLA-I substantially reduces the load and diversity of the non-self peptidome susceptible to be presented by the HLA-I^12, 14^. However, the absolute loss of potential neoepitopes depends on the specific HLA-I allotypes and the total burden of somatic alterations. This prompted us to analyze whether LOH of HLA-I tends to involve the loss of the allele/s with the highest neoepitope ratio (i.e., higher number of predicted neoepitopes compared to the alternative allele; see Fig. 4a, methods Tumor specific neoepitopes and Supp. Note 2). We observed a positive association between the neoepitope ratio and the frequency of the allele with higher neoepitope repertoire to be lost across both the primary and metastatic datasets (Fig. 4b and Fig. 4e). This trend was significantly different from a neutral scenario where both alleles are equally likely to be lost independently of their neoepitope repertoire (Kolmogorov–Smirnov test metastatic p-value=2.47E-5 and primary p-value=1.26E-7; methods calculation and randomization neoepitope ratio). The association between neoepitope ratio and the loss frequency became stronger when selecting for focal and highly focal LOHA HLA-I events (p-value=3.26E-9 and p-value=1.65E-10 for metastatic and primary respectively). However, it was indistinguishable from a neutral scenario for non-focal LOH-HLA (p-value=0.24 and p-value=0.11 for metastatic and primary respectively), showing that non-focal LOH HLA-I does not select for the allele with the highest neoepitope repertoire and that its high recurrency in certain cancer types may be associated with other selective forces.

**Figure 4.**
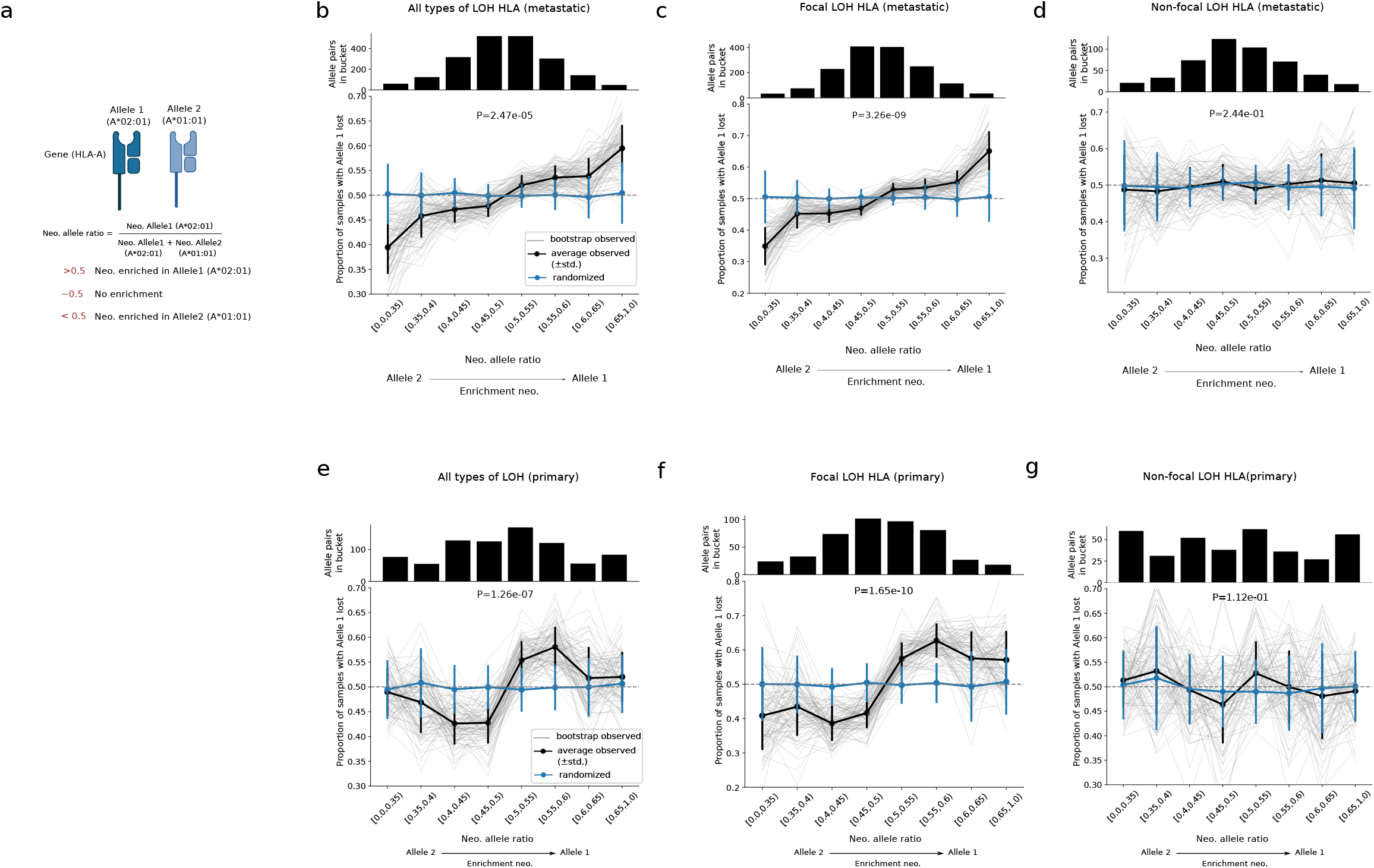
LOH HLA-I and the neoepitope load. a) Visual depiction of the neoepitope allele ratio and its significance. b) top panel, number of allele pairs in each neoepitope allele ratio bucket in metastatic samples harboring LOH of HLA-I. bottom, representation of the mean observed (black) and randomized (blue) neoepitope allele ratio across 100 bootstraps (thicker lines). Vertical error bars represent the standard deviation of the neoepitope allele ratio. Narrow black lines represent the observed neoepitope allele ratio values across the 100 bootstraps. P-values were calculated using the Kolmogorov–Smirnov test. c) and d) are identical but subsampling for focal and non focal LOH of HLA, respectively. e), f) and g) equivalent panels for the primary dataset. neo., neoepitope.

### Positive selection of GIE alterations beyond HLA-I

Alterations in other pathways beyond the HLA-I locus may also lead to immune surveillance escape. Hence, we explored signals of positive selection across 18 genes associated with five immune escape pathways (pathways 2-6 in Fig. 1a). Grouped pan-cancer analysis of the dN/dS ratio in these pathways (covering a total of sixteen genes, excluding those whose oncogenic mechanism is based on copy number amplification, see methods Positive selection somatic mutations and indels) revealed a greater than one ratio for nonsense, splice site and truncating variants in both the metastatic and primary datasets (Fig. 5a), which was indicative of positive selection. We next performed a gene and cancer type specific positive selection of genes involved in antigen presentation machinery, IFN-γ pathway, PD-L1, CD58 and epigenetic immune escape (methods positive selection and Supp. Table 5). We found at least two genes with signals of positive selection involved in the antigen presentation pathway: B2M and CALR. B2M was considered as significantly mutated (dNdScv q-value<0.1) in colorectal, kidney clear carcinoma, non-small cell lung cancer and diffuse large B-cell lymphoma and the two pan-cancer groups (Fig. 5b and Fig. 5c). Focal B2M loss was significantly recurrent in ovarian cancer, skin melanoma and diffuse large B-cell lymphoma as well as the pan-cancer metastatic cohort (G-test goodness of fit q-value<0.1; Fig. 5d). Similarly, CALR was targeted by focal recurrent biallelic loss in primary skin melanoma. Higher than expected frequency of focal biallelic deletions of several IFN-γ pathway genes, including JAK1, JAK2 and IRF2 was also observed. CD58 harbored higher than expected number of non-synonymous mutations and homozygous deletions in diffuse large B-cell lymphoma and in the pan-cancer metastatic group. Finally, the chromatin modifier SETDB1 harbored highly-focal copy number amplifications in multiple cancer types, such as metastatic non-small cell lung cancer (Fig. 5e) and primary breast cancer, among others. In summary, immune escape alterations disrupting several immune related pathways, other than the HLA-I, are recurrently exploited by tumors to hamper the presentation and recognition of non-self antigens.

**Figure 5.**
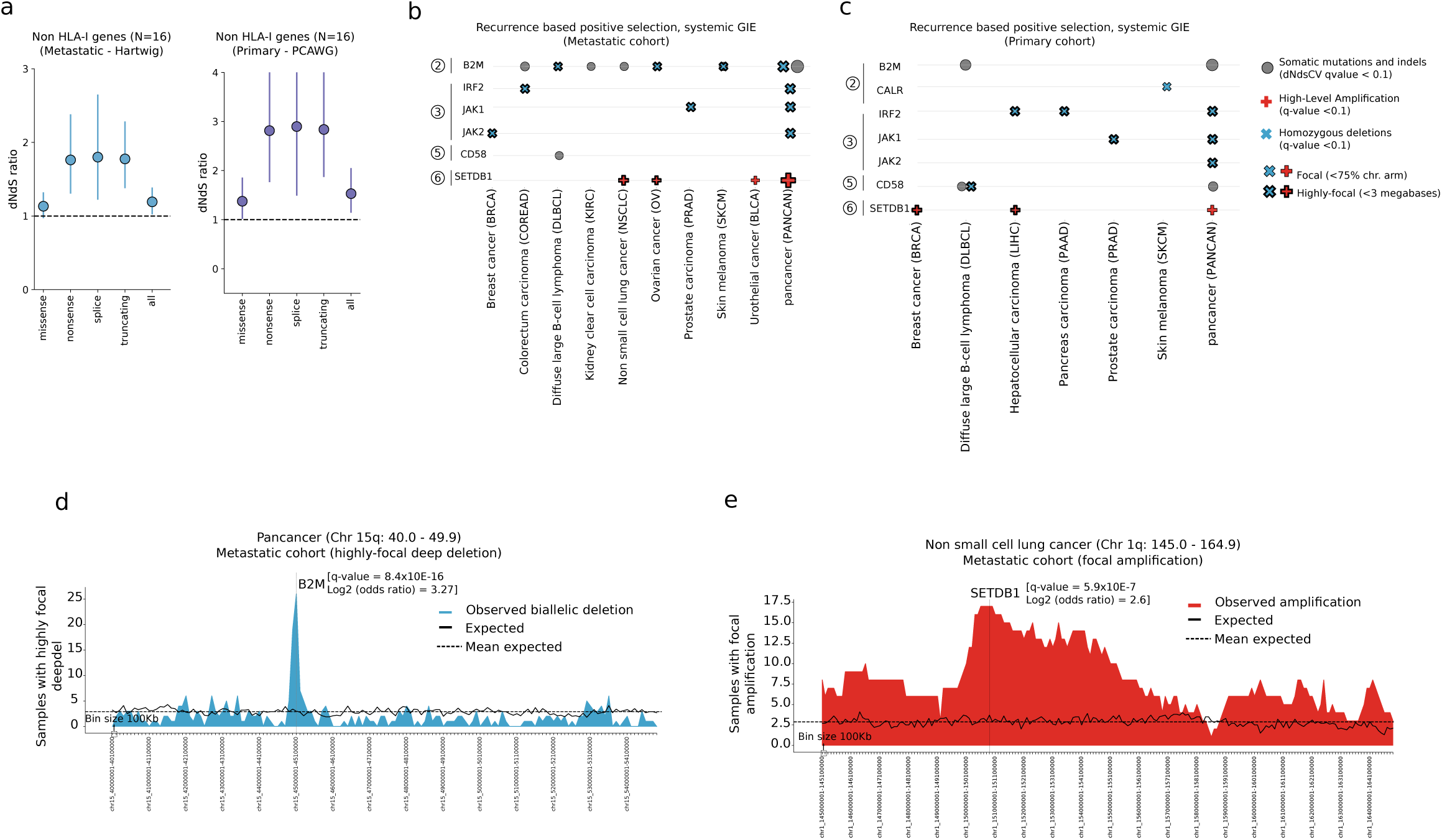
Positive selection of systemic HLA-I alterations. a) Pan-cancer dN/dS ratios of non HLA-I genes in the metastatic (left) and primary (right) datasets. Vertical lines represent the 5% and 95% confidence intervals. b) Representation of gene and cancer type specific positive selection of GIE systemic genes in the metastatic and c) primary cohorts. The pathway number attributed to each gene is displayed next to the gene name (relative to Fig. 1a). d) Distribution of highly-focal biallelic deletions surrounding the B2M gene, spanning from chr15:40.0Mb to chr15:49.9Mb in the pan-cancer metastatic cohort. e) Distribution of highly-focal copy number amplification surrounding the SETDB1 gene, spanning from chr1:145.0Mb to chr1:164.9Mb in the metastatic non-small cell lung cancer (NSCLC) cohort. Each bin represents 100Kbs. Dashed horizontal lines represent the expected mean after randomization. Vertical dashed lines highlight the gene genomic location. Mb, megabase. Kbs, kilobases.

### GIE association with cancer genomic features

We next explored whether, aside from cancer type intrinsic differences, there were other cancer genomic features linked to an increase or decrease in GIE prevalence. We performed a cancer type specific univariate logistic regression to screen the association of 95 genomic features, such as the mutation burden per mutation type, DNA repair deficiency status or the presence of HLA-I supertypes; across 32 cancer types (see methods tumor genomic features and GIE risk and Supp. Table 6). Overall, 46 genomic features showed a statistically significant association with GIE in at least one cancer type (Fig. 6a). As expected, TMB and patient’s neoepitope load, defined as the collection of the HLA-I germline allele specific neoepitopes, were strongly associated with GIE events across multiple cancer types (logistic regression corrected p-value<0.05 and Log_2_(Odds Ratio)>0.0; Fig. 6a and Supp. Fig. 4a-c). Interestingly, clonal TMB and clonal neoepitope load showed a strong positive association whereas neither subclonal TMB or subclonal neoepitope load were predictive of GIE in any of the cancer types (Fig. 6a-c), highlighting the relevance of mutation cellularity in triggering immune responses. Gene fusions and structural variants may also be the source of non-self antigens. In fact, in our dataset, these features and fusion derived neoepitopes were significantly associated with GIE in diffuse large B-cell lymphoma, breast, ovarian and non-small cell lung cancer (Fig. 6a and Supp. Fig. 4a), which emphasizes the usefulness of considering non-canonical sources of neoepitopes beyond small non-synonymous variants in coding regions.

**Figure 6.**
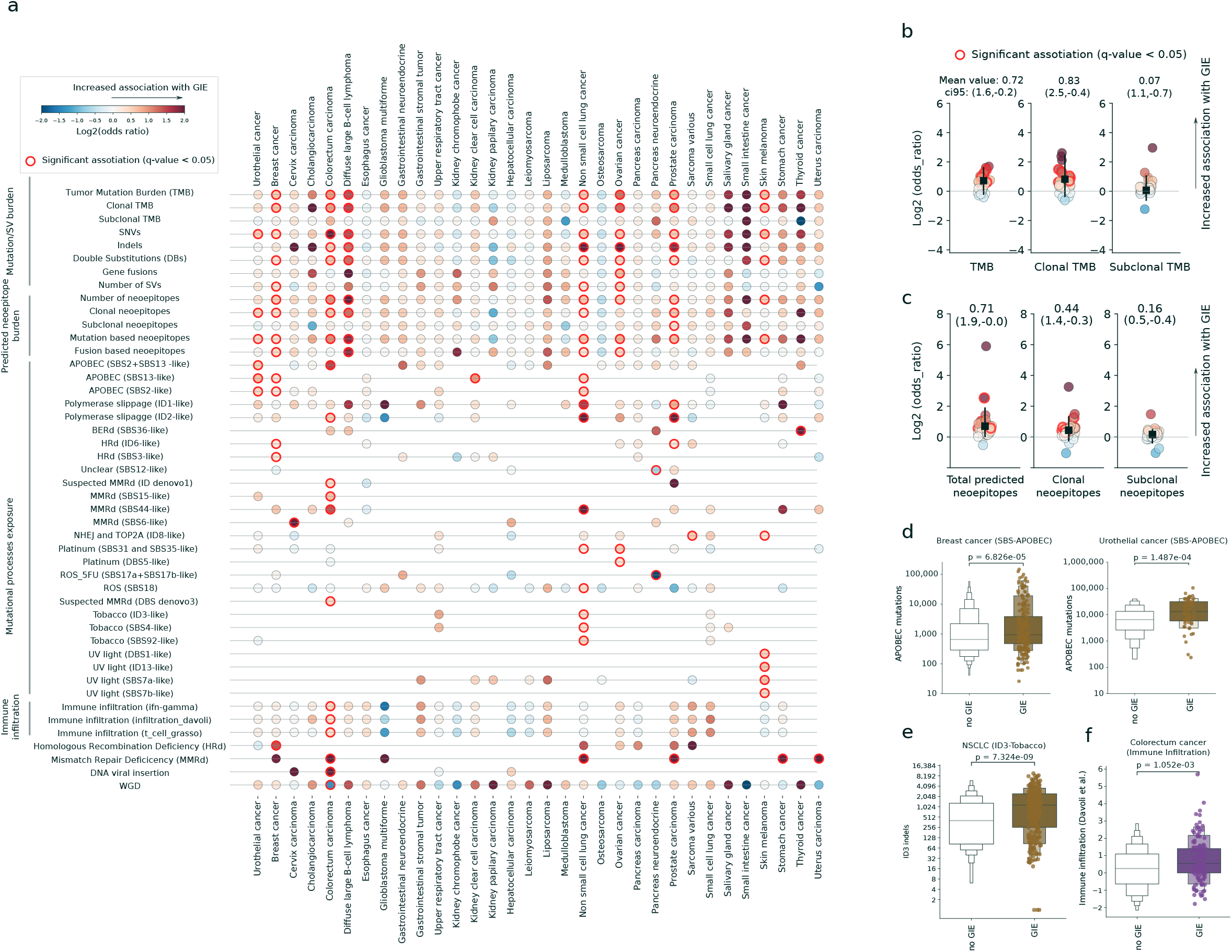
GIE association with cancer genomic features. a) Heatmap displaying the association of 46 genomic features with GIE frequency across 32 cancer types. Features displayed have, at least, one significant cancer type association with GIE alterations. Significant associations are highlighted by a red border line. Dot colors are coloured according to the log_2_(odds ratio). b) from left to the right, dotplot representations of the TMB, clonal TMB and subclonal TMB log_2_(odds ratio) across the 32 cancer types. Black square represents the average values. The error bar represents the 95% and 5% confidence intervals (ci95). Horizontal lines represent a neutral scenario with log2(odds ratio) equal to 0. c) analogous representation for predicted neoepitopes, clonal neoepitopes and subclonal neoepitopes, respectively. d) Comparison of the APOBEC mutational exposure between samples bearing GIE alterations (GIE) and wild-type (no GIE) in breast cancer (left) and urothelial cancer (right). e) Similar comparison for Tobacco COSMIC ID3-like indels load in non-small cell lung cancer (NSCLC). f) Comparison of immune infiltration estimates^38^ between samples bearing GIE alterations (GIE) and wild-type (no GIE) in colorectal cancer. Box-plots: center line, median; box limits, first and third quartiles; whiskers, lowest/highest data points at first quartile minus/plus 1.5× IQR. P-values are calculated using a two-sided Mann–Whitney U test.

Exposure to certain endogenous and exogenous mutational processes have been correlated with increased immunogenicity^27^ and response to ICI^28, 29^. We therefore performed a cancer type specific and molecular age controlled logistic regression of mutational processes exposure to assess their association with GIE risk (see methods mutational signatures). DNA repair deficiency mutational signatures, including Mismatch Repair Deficiency (MMRd, SBS6, SBS16, SBS44, ID1, ID2, ID denovo1 and DBS denovo3), Base Excision Repair deficiency (BERd, SBS36 see Supp. Fig. 4d) and Homologous Recombination deficiency (HRd, SBS3 and ID6) were broadly associated with increased GIE incidence (logistic regression corrected p-value<0.05 and Log_2_(Odds Ratio)>0.0; Fig. 6a and Supp. Fig. 4a). Sample specific HRd and MMRd binary classification provided consistent results (Fisher’s exact test adjusted p-value<0.05). Remarkably, exposure to the APOBEC family of cytidine deaminases (SBS2 and SBS13) was strongly associated with GIE in multiple cancer types, including breast and urothelial cancer (Fig. 6d), among others. Exposure to several exogenous mutational processes also exhibited a significant association with increased GIE risk. Concretely, tobacco exposure (SBS4, SBS92 and ID3; Fig. 6e) in non-small cell lung cancer (NSCLC), UV light mutation burden (SBS7a, SBS7b, DBS1 and ID13; Supp. Fig 6d) in skin melanoma and platinum treatment mutation load (SBS31-SBS35-like and DBS5) in ovarian (Supp. Fig 6e) and NSCLC were significantly linked to an increased incidence of GIE events. Lastly, other mutational signatures resulted in a significant association (Fig 6a.), although their etiology and role in immune evasion is less apparent.

Other tumor genomic features were also correlated with GIE. More specifically, in colorectal cancers, which also include some anal cancer patients, HPV viral integration was positively associated with GIE, suggesting that GIE provides a selective advantage in tumor cells harbouring viral DNA (Fig. 6a and Supp. Fig. 4a). Most of HPV+ colorectum samples are likely anal cancer classified as colorectum due to variation in how detailed primary tumor classes were registered in the studies in which the metastatic samples were collected. Similarly, we also found a negative association between whole genome duplication (WGD) and GIE, likely explained by diploid karyotype of hypermutated and microsatellite unstable tumors, often harboring systemic immune escape alterations^30^. Finally, high-immune infiltration as determined by several RNA-Seq-based deconvolution measurements for those samples for which such data was available (see methods of immune infiltration deconvolution) was significantly linked with higher GIE incidence in colorectal carcinoma (Fig. 6a, Fig. 6f and Supp. Fig. 4a), which was in agreement with previous reports^4^. Finally, other factors, such as the HLA-I supertype, the germline HLA-I divergence or exposure to previous treatments failed to attain significant association with GIE. All the screened molecular features alongside their cancer type specific significance coefficients are available at Supp. Table 6.

In conclusion, the presence of GIE is strongly associated with certain tumor genomic features, including high TMB, exposure to certain endogenous and exogenous mutagenic mutational processes or the presence of viral DNA, and can not be solely explained by cancer type intrinsic differences.

### Tumors tailor their immune evasion strategy depending on the TMB

An increase in mutation load leads to the generation of neoepitopes susceptible to be recognized as neoantigens by the T-Cell receptors (TCRs) of surrounding T-cells. Consequently, an increase in TMB may require immune escape alterations to prevent clearance by the immune system (see above). We examined the frequency of GIE alterations in tumors across 20 evenly distributed TMB buckets (see methods GIE and TMB dependence). We found that GIE frequency steadily increased with increasing TMB (Fig. 7a). Interestingly, in the bucket grouping samples with ∼10-13 muts/Mb, a threshold frequently used as response to ICI, we observed an average GIE frequency of 0.32 ±0.03 std. Similarly, in the group of samples between 26-36 muts/Mb, mostly including hypermutated tumors, the average frequency was 0.49 ±0.1 std., while beyond ∼95 muts/Mb (considered ultra-hypermutated tumors^31^) we identified GIE alterations in more than 70% of samples (>0.72 ±0.06 std.). Moreover, using the burden of predicted neoepitopes based on the germline HLA-I profile as baseline revealed a near-uniformly increasing distribution across the neoepitope buckets, which becomes sharper after the 17th bucket (i.e., number of neoepitopes greater than 1,410 and lower than 1,937, see Fig. 7b).

**Figure 7.**
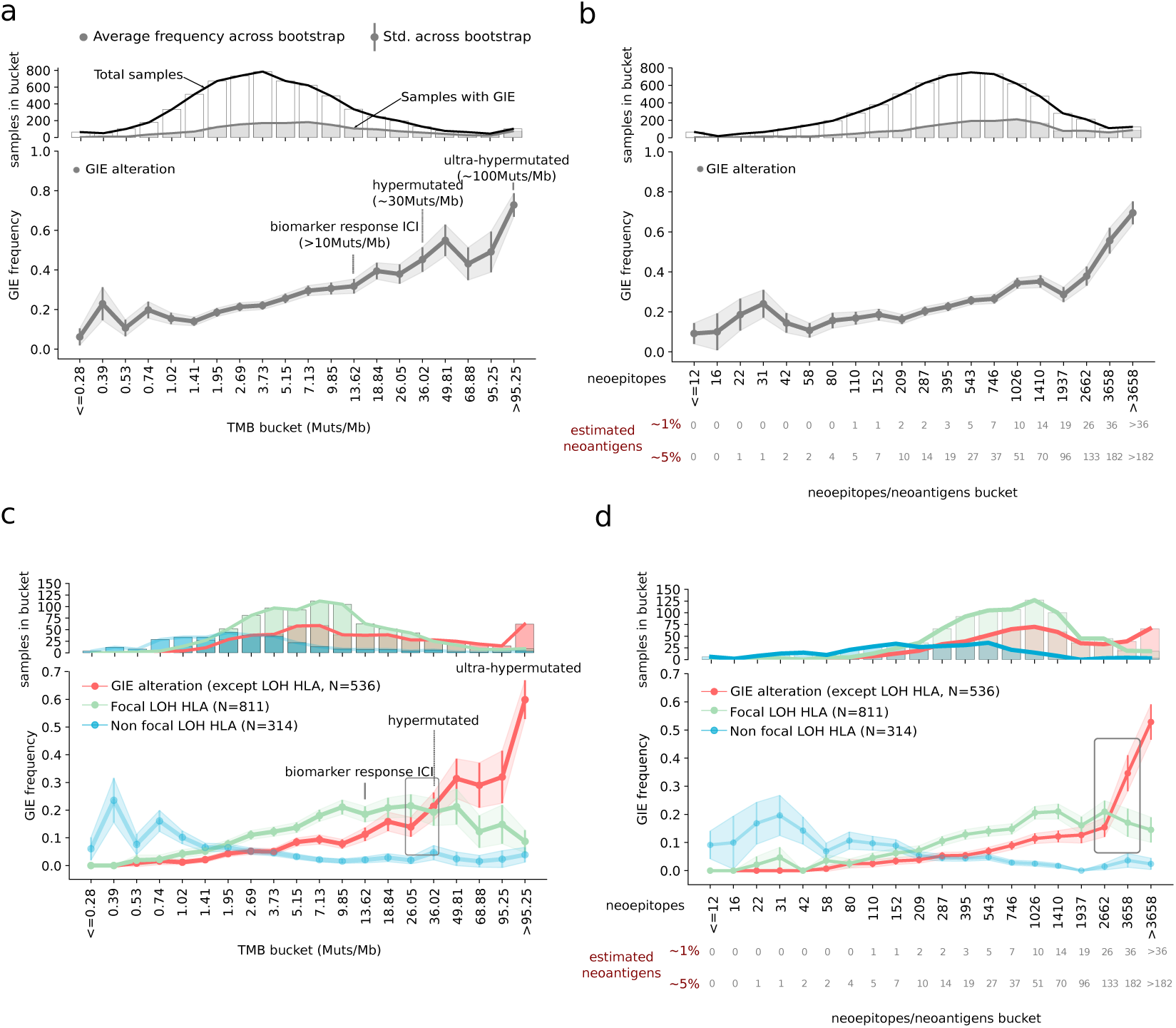
Immune evasion mechanism and TMB. a) Top panel, number of total (white bars with a black contouring line) and GIE-present (gray bars with gray contouring line) samples across the TMB buckets. bottom panel, representation of GIE frequency across twenty evenly distributed TMB buckets. Gray dots represent the average GIE frequency across 1,000 bootstraps. Error bars and the gray shade represent ± standard deviation. b) Using predicted neoepitopes as baseline buckets. Bottom labels, number of estimated neoantigens as a relative percentage (1% and 5%) of the number of predicted neoepitopes in the bucket. c) and d) related to a) and b) respectively, but splitting by type of GIE alterations. Inner boxes highlight the bucket where non-LOH HLA-I frequency (red) surpasses focal LOH HLA-I (green). Muts/Mb, mutations per megabase.

GIE involves a diverse collection of immune escape pathways, which encompasses HLA-I targeted and systemic tumor specific alterations. We then analyzed the relationship between the tumor mutation burden and the presence of specific GIE alterations across the six immune escape pathways included in this study (see Fig. 1a). Overall, the observed frequency distributions across these pathways were remarkably different (Supp. Fig. 5a). In fact, different types of HLA-I alterations showed a distinctive frequency distribution along the TMB buckets. Non-focal LOH of HLA-I were primarily present in low-TMB tumors, supporting the rationale that it is likely a passenger event. Conversely, focal LOH of HLA-I showed a clear enrichment for mid and high TMB tumors peaking around ∼10 muts/Mb (i.e., 9.85 muts/Mb bucket with average frequency of 0.19 ±0.03 std.) and displaying an inverted U-shaped distribution. Finally, mutations in HLA-I genes were more frequent in hypermutated and ultra-hypermutated tumors (i.e., from the ∼26-36 muts/Mb bucket onwards). Similarly, alterations in the antigen presentation machinery and in the IFN-γ pathway (pathways number 2 and 3, respectively) were predominantly found in high and very high TMB tumors, becoming the dominant GIE mechanism in hypermutated tumors (Supp. Fig 5a). The remaining pathways (CD58, PD-L1 and epigenetic remodeling by SETDB1) did not show any clear TMB preference, likely due the lower prevalence of these alterations in our dataset. Taken together, our results reveal a non-overlapping pattern where non-focal LOH of HLA is primarily observed in low TMB tumors, focal LOH of HLA in mid-high TMB tumors and other GIE alterations (mainly driven by the inactivation of the antigen presentation machinery and the IFN-γ pathways as well as loss-of-function mutations in HLA-I) dominating in hypermutated patient samples (Fig. 7c).

Using the number of predicted neoepitopes (derived from point mutations, indels and gene fusions) instead of TMB as baseline revealed consistent distributions (Supp. Fig. 5b and Fig. 7d). While focal LOH of HLA-I was the most frequent GIE mechanism in tumor samples with ∼200-2,600 predicted neoepitopes (corresponding to 2-133 estimated neoantigens^6, 32^) the other GIE alterations group became the most frequent GIE event after the 2,662 neoepitopes bucket (26-133 neoantigens). In conclusion, tumors select their GIE mechanism depending on the neoepitope burden, where mid-high TMB tumors primarily leverage LOH of HLA-I and hypermutated tumors tend to rely on systemic immune evasion as the preferred GIE mechanism.

## Discussion

A hallmark of tumorigenesis is the capacity of cancer cells to evade immune system surveillance^1^. To decipher the landscape of genetic immune escape alterations, we have analyzed their prevalence and impact across six major pathways in thousands of uniformly processed primary and metastatic tumors from 58 cancer types. We addressed the complexity of identifying tumor-specific HLA-I alterations by developing LILAC, a tool that performs a robust characterization of the HLA-I locus in tumor samples. In fact, LILAC showed a near-perfect HLA typing performance on WGS data, highlighting its potential for use in routine diagnostics.

Combining LILAC with a universal pipeline to identify somatic alterations provided a patient level annotation of GIE events. On average, one in four patients bore GIE events, primarily as a result of LOH of HLA-I and displaying a highly variable prevalence across cancer types. Despite the high frequency of LOH of HLA-I, biallelic deletion of both HLA alleles was an extremely unusual event, suggesting the importance of expressing a minimal amount of HLA-I molecules to avoid immune-alerter signals^33^.

Remarkably, our results also showed that the frequency of genetic immune escape alterations in metastatic patients are comparable to their primary counterparts across most of the cancer types, suggesting that early stages of tumorigenesis have already acquired the capacity to escape from immune system recognition.

We also observed that immune escape alterations are positively selected during tumor evolution. Loss-of-function mutations in HLA-A and HLA-B were recurring across several cancer types, which is in agreement with the tumor suppressor role of these genes. Nevertheless, HLA-C did not show a significant enrichment in inactivating variants which may imply that its expression is needed to avoid NK-mediated immunity^34^ and that the neoepitope repertoire of this gene is generally lower compared to HLA-A and HLA-B. LOH of HLA I was observed more frequently than expected across multiple cancer types. Moreover, focal LOH of HLA-I events primarily targeted the HLA allele with the highest neoepitope repertoire whereas non-focal LOH of HLA-I did not. While we can not rule out the possibility that the loss of several HLA-I alleles provides an accessory fitness gain, non-focal LOH of HLA-I does not seem to be the primary target of these large-scale events. We also observed that genes involved in pathways such as the antigen presentation machinery or the IFN-γ signaling were also recurrently mutated in multiple cancer types, indicating that tumors harness multiple molecular pathways to circumvent the selective pressure imposed by the immune system.

Our results showed that both tumor type and patient-specific features contribute to the presence and diversity of GIE alterations. Several factors such as the amount of small and structural somatic variants (TMB), exposure to certain mutational processes or HPV DNA integration are significantly associated with an increased GIE prevalence, revealing the potential role of these features in raising the visibility to the immune system. While the association with an increased immunogenicity of certain genomic features such as the total TMB or the global neoepitope load is evident, others such as the presence of particular mutational processes require further studies. Moreover, patient specific integration of immunogenic features alongside GIE alterations could potentially improve current biomarkers of response to immunotherapy treatment.

Finally, we observed that tumors leveraged different GIE strategies according to the total TMB and the global neoepitope load. While LOH of HLA-I was the most frequent mechanism in mid and high TMB tumors, loss of certain HLA-I alleles was apparently not sufficient to cope with the elevated neoepitope load of hypermutated and ultra-hypermutated tumors, where a systemic GIE mechanism, such as antigen presentation abrogation or IFN-γ inactivation, is therefore needed.

This study considered a collection of highly-confident and well characterized genetic immune escape alterations across six well characterized immune related pathways. However, there are likely other potential sources of genetic immune evasion, which include not only alternative molecular pathways such as the HLA-II^35^ and the killer immunoglobulin-like receptor^33^ (KIR); but also other types of alterations such as germline variants^36^. Additionally, recent studies have shown the role of epigenetic modifications in modulating tumor immunogenicity^2, 37^. Hence, the combination of somatic and germline alterations with other -*omics* such as epigenomics, proteomics, single-cell RNA sequencing or TCR-Seq will provide a more thorough picture of the interplay between tumor (epi)genomic alterations and the immune system.

In conclusion, our study provided a comprehensive landscape of immune escape alterations across primary and metastatic tumors. Our results suggest that the diversity and frequency of immune escape mechanisms are highly shaped by the cancer type and by certain tumor genomic features and that the metastatic bottleneck does not generally introduce significant changes in this landscape. Further studies will be needed to disclose how these findings should be harnessed in therapeutic interventions.

## Methods

### Data collection and processing

#### Hartwig metastatic cohort

The Hartwig Medical Foundation sequences and characterizes the genomic landscape for a large number of metastatic patients across the Netherlands. A detailed description of the consortium and the whole patient cohort has been described in detail in Priestley et al^18^. In this study, the Hartwig cohort included 4,784 metastatic tumor samples from 4,468 patients.

The Hartwig patient samples have been processed using the Hartwig analytical pipeline5 (https://github.com/hartwigmedical/pipeline5) implemented in Platinum (https://github.com/hartwigmedical/platinum). Briefly, Platinum is an open-source pipeline designed for analyzing WGS tumor data. It enables a comprehensive characterization of tumor WGS samples (e.g., somatic point mutations and indels, structural variants, copy number changes, etc.) in one single run. See variant calling section for further information.

Hartwig samples that failed to provide a successful pipeline output, potential non-tumor samples, with purity lower than 0.2, with TMB <50 SNVs/indels, lacking sufficient informed consent for this study or without enough read coverage to perform 4 digit HLA typing (see below, LILAC section) were discarded. Similarly, for patients with multiple biopsies we selected the tumor sample with the most recent biopsy date, and if this information did not exist we selected the sample with the highest tumor purity. However, some Hartwig patients had biopsies from different primary tumor locations. In these cases, we kept at least one sample from each primary tumor location, and when there were multiple samples from the same primary tumor location, we applied the aforementioned biopsy date and tumor purity filtering criteria. A total number of 4,514 Hartwig samples were whitelisted and used in this study (Supp. Fig. 2a and Supp. Table 3).

Pre-processed RNA-seq data by ISOFOX (https://github.com/hartwigmedical/hmftools/tree/master/isofox) was available for 1,864 Hartwig samples and was consequently used in the immune infiltration deconvolution analysis. Patient clinical data was obtained from the Hartwig database. Cancer types labels were harmonized to maximize the number of samples with comparable tumor types with the PCAWG dataset (see Supp. Table 3).

#### PCAWG primary cohort

The PCAWG cohort consisted of 2,835 patient tumors, and access for raw sequencing data for the PCAWG-US was approved by National Institutes of Health (NIH) for the dataset General Research Use in The Cancer Genome Atlas (TCGA) and downloaded via dbGAP download portal. Raw sequencing access to the non-US PCAWG samples was granted via the Data Access Compliance Office (DACO). A detailed description of the consortium and the whole patient cohort has been described in ref^19^.

The samples were fully processed using the same cancer analytical pipeline applied to the Hartwig cohort. This enabled a harmonized analysis and eliminated the potential biases introduced by applying different methodological approaches. Samples that failed to provide a successful pipeline output, with a tumor purity below 0.2, potential non-tumor samples, blacklisted by the PCAWG original publication^19^ or without enough read coverage to perform 2-field HLA typing were discarded. Similarly, for patients with multiple samples, we selected the first according to the aliquout_id alphabetical order. A total number of 1,943 were whitelisted and used in this study (Supp. Fig. 2a and Supp. Table 3).

Pre-processed gene level expression data was downloaded for 1,118 samples from the ICGC portal (https://dcc.icgc.org/releases/PCAWG/transcriptome/gene_expression/_star_fpkm_uq. <u>v2_aliquot_gl.tsv.gz</u>). ENSEMBL identifiers were mapped to HUGO symbols. 949 of these samples belonged to biopsies selected for this study and were therefore used for the RNA analyses in PCAWG samples.

The most recent clinical data was downloaded from the PCAWG release page (https://dcc.icgc.org/releases/PCAWG/) on August 2021. Cancer types labels were harmonized to maximize the number of samples with comparable tumor types with the Hartwig dataset (see Supp. Table 3).

#### Variant calling

As mentioned above, the Hartwig and PCAWG samples have been uniformly processed using the Hartwig analytical pipeline5 (https://github.com/hartwigmedical/pipeline5) implemented in Platinum (https://github.com/hartwigmedical/platinum). Briefly, sequencing reads were mapped to GRCh37 using BWA (v0.7.17). GATK (v3.8.0) Haplotype Caller was used for calling germline variants in the reference sample. SAGE (https://github.com/hartwigmedical/hmftools/tree/master/sage, v2.2) was used to call somatic single and multi nucleotide variants as well as indels. GRIDSS^39^ (v2.9.3) was used to call simple and complex structural variants. PURPLE (https://github.com/hartwigmedical/hmftools/tree/master/purple) combines B-allele frequency (BAF) from AMBER (https://github.com/hartwigmedical/hmftools/tree/master/amber, v3.3), read depth ratios from COBALT (https://github.com/hartwigmedical/hmftools/tree/master/cobalt, v1.7), and structural variants from GRIDSS to estimate copy number profiles, variant allele frequency (VAF) and variant clonality. LINX (https://github.com/hartwigmedical/hmftools/tree/master/linx) interprets structural variants (to identify simple and complex structural events) from PURPLE, and also detects gene fusions, viral DNA integrations, and homozygously disrupted genes. Importantly, we ensured that mutation (simple and complex) filtering and annotation tools were run with the same versioning for PCAWG and the Hartwig cohorts. For PURPLE we relied on v2.53 whereas for LINX we used v1.16.

### LILAC

#### Overview

LILAC is a WGS framework to determine the HLA class I types for the germline of each patient as well as determining the status of each of those alleles in the tumor including complete loss of one or more alleles, allele specific somatic mutations and allelic imbalance.

LILAC provides several conceptual and practical improvements over the numerous available tools for HLA-I typing: i) increased accuracy for WGS samples with coverage between 30-100X, particularly remarkable for rare alleles, ii) integrated analysis of paired tumor-normal sample data to call allele-specific copy number and assignment of somatic variants to specific alleles, iii) detection of novel germline variants and/or alleles (including indels) via analysis of unmatched fragments iv) full report of quality control metrics and number of fragments assigned to each HLA-I allele and, finally, v) identification of HLA-Y presence (a pseudogene with high similarity to HLA-A present in up to 20% of the population but is not present in the reference genome).

LILAC relies on the somatic point mutations and copy number estimations to estimate the tumor HLA-I status. Lastly, LILAC supports GRCH37, hg19 and hg38 (with no alt) reference genomes (Fig. 1b and Supp. Note 1). LILAC is avaliable at https://github.com/hartwigmedical/hmftools/tree/master/lilac.

#### Algorithm

The starting point for the LILAC algorithm is the complete set of possible 4 digit alleles and all the fragments aligned to HLA-A, HLA-B and HLA-C. LILAC algorithm begins with collecting all fragments which are not duplicates and have:

- At least one read with an alignment overlapping a coding base of HLA-A, HLA-B or HLA-C; and
- all alignments within 1000 bases of a HLA coding region; and
- a mapping quality of at least 1

The algorithm then has 2 main phases to determine the germline alleles: an ‘elimination’ phase which aims to remove allele candidates that are clearly not present and an ‘evidence’ phase where LILAC considers all possible sets of 6 alleles amongst the remaining candidates and chooses the solution that best explains the fragments observed.

After the germline alleles are determined, LILAC determines the tumor copy number and any somatic mutations in each allele. Note that if more than 300 bases of the HLA-A,HLA-B and HLA-C coding regions have less than 10 coverage, then LILAC will fail with errors and will not try to fit the sample.

Supp. Note 1 presents full details of LILAC’s algorithm and its implementation.

#### Germline-tumor agreement comparison

We assessed LILAC’s robustness to perform HLA-I typing compared to two state-of-the art tools: Polysolver^15^ (v4, *reference genome* hg19, *ethnicity* Unknown, *insertCalc* 0 and *includeFreq* 0) and xHLA^20^ (with default parameters). We first retrieved GRCh37 aligned reads including the MHC-I locus (chr6:29,854,528-32,726,735) for the PCAWG and Hartwig germline and tumor samples. For each of these three tools, we first independently run the HLA-I typing for the germline and for the tumor across all available samples (see above, Hartwig and PCAWG cohort), and then we annotated whether there was a perfect agreement between them based on the inferred 2-field HLA-I haplotypes. Samples that failed to provide an output by any of the three methods were not included in the comparison. A total of 4,774 Hartwig and 2,099 successfully provided germline and tumor HLA haplotypes and were used in this analysis. Moreover, to reduce potential effects of tumor specific alterations on the HLA-I genes, we performed a similar comparison but limited to samples without HLA-I alterations (i.e. somatic mutations or LOH HLA events) according to LILAC.

#### Crosswise tools HLA-I haplotype comparison

We also assessed LILAC’s agreement with two widely used tools for HLA typing: Polysolver (v4, *reference genome* hg19, *ethnicity* Unknown, *insertCalc* 0 and *includeFreq* 0) and xHLA (with default parameters). For each of these tools we first ran the germline HLA typing and we then performed the comparison based on the four digit HLA-I type annotation. Samples that failed to provide an output by any of the three methods were not included in the comparison. A total of 4,774 Hartwig and 2,099 successfully provided germline and tumor HLA haplotypes and were used in this analysis. We only considered that two, or three respectively, tools have an agreement if the four digit HLA-I alleles perfectly match among them.

#### Copy number estimation of HLA-I compared to other genes

We aimed to evaluate whether the polymorphic nature of the HLA-I locus could have a negative impact on the tumor copy number estimation and subsequent annotation of HLA-I individual alleles. A proxy for incorrect tumor copy number estimation is the difficulty to assign an integer copy number. Therefore, we compared the proportion of samples with HLA-I genes with a purity adjusted integer copy number (i.e., estimated minor and major allele copy number <= 0.3 or >= 0.7) compared to other 1,000 randomly selected genes across the human exome. We performed this comparison across the two cohorts used in this study. Only samples with sufficient quality according to LILAC were used in this comparison.

#### Experimental validation by high-to allelic resolution HLA typing

We selected 96 Hartwig samples (10 from the tumor and 86 from the germline) to assess LILAC’s agreement with an orthogonal approach based on high-to allelic resolution HLA typing (see below). The selection of samples were prioritized based on the following criteria: i) sample availability, ii) disagreement of LILAC with either xHLA or Polysolver, iii) challenging cases due to presence of rare alleles and iv) tumor samples bearing either somatic mutations or LOH of HLA-I. One germline sample failed to provide output with sufficient quality and was therefore not included in the final comparison (see Supp. Table 2).

HLA genes were amplified with the NGSgo® MX11-3 (GenDx) amplification strategy and libraries were prepared using the NGSgo® Library Full Kit (GenDx); both according to manufacturer’s instructions. Libraries were sequenced on the MiSeq (Illumina) and the generated .fastq files were analyzed using the HLA typing analysis software NGSengine® (GenDx), 2.24.0, using the IMGT3.44 reference database. All data was reviewed by two independent reviewers and all exon heterozygous positions deviating from standard patterns were inspected and interpreted manually.

#### Tumor-normal paired mode in Hartwig and PCAWG

All pre-selected PCAWG and Hartwig samples (see above and Supp. Fig. 2a) were then processed by LILAC using the tumor-normal pair mode, which relied on the germline and tumor raw HLA-I files, the somatic mutation calls by SAGE (https://github.com/hartwigmedical/hmftools/tree/master/sage) and the copy number estimations by PURPLE (https://github.com/hartwigmedical/hmftools/blob/master/purple/) (both outputs were available as part of the Platinum Hartwig pipeline5 output).

All the pre-selected Hartwig samples but one (4,514 samples out of 4,515) were successfully processed by LILAC whereas 1,943 out of 2,359 PCWAG samples successfully achieved LILAC’s quality control criteria. The main failure reason of PCAWG samples was insufficient coverage to perform a four digit HLA-I typing, and therefore they can not be further considered in this study.

As a result of the pipeline we obtained 2-field HLA-I class types for all successful samples alongside annotation of somatic mutations and the copy number estimations mapping for each allele.

### GIE alterations

#### Definition

We searched in the literature for somatic genomic alterations that are robustly and recurrently associated with immune evasion. We stratified the reported alterations into six major pathways (Fig. 1a and Supp. Table 1):

1. **The HLA-I**. Somatic alterations in the HLA-A, HLA-B and HLA-C genes have been extensively reported as a mechanism for immune evasion across several cancer types^11, 12, 14, 25^. We considered LOH HLA, homozygous deletions and somatic non-synonymous mutations on these genes as immune evasion alterations. We defined LOH for HLA-A, HLA-B and HLA-C as those cases with a minor allele ploidy below 0.3 and a major allele ploidy greater than 0.7 according to LILAC annotation. We also relied on LILAC mapping of somatic mutations into HLA-A, HLA-B and HLA-C alleles to report samples with non-synonymous alterations. Finally, we also used LILAC allele specific tumor copy number estimations to annotate samples with homozygous deletions of HLA-A/B/C genes. A gene was homozygously deleted in a sample if the estimated minimum tumor copy number of the gene was lower than 0.5.
2. **The antigen presentation pathway.** Several studies have reported the immunomodulatory effect of somatic inactivation of genes involved in the antigen presentation machinery (see Supp. Table 1 for gene-specific references). The most recurrent alteration is B2M inactivation, but there are other genes involved in antigen presentation and antigen presentation activation, whose inactivation has been linked to increased immune evasion, including CALR, TAP1, TAP2, TAPBP, NLRC5, CIITA and RFX5. We defined inactivation events as mono-allelic and bi-allelic clonal loss of function mutations (frameshift variant, stop gained, stop lost, splice acceptor variant, splice donor variant, splice region variant and start lost), bi-allelic clonal non-synonymous mutations not included in the former group (e.g. missense mutations) and homozygous deletions. A gene was homozygously deleted in a sample if the estimated minimum tumor copy number of the gene was lower than 0.5.
3. **The IFN-γ pathway.** IFN-γ is a cytokine with known pro-apoptotic and immune booster capacities. Hence, it has been reported that tumors frequently leverage somatic alterations targeting IFN-γ receptors and downstream effectors to evade immune system surveillance (see Supp. Table 1 for gene-specific references). More specifically, we considered inactivation events (see above for specifics of which type of alterations are included) in JAK1, JAK2, IRF2, IFNGR1, IFNGR2, APLNR and STAT1 have been probed to have the ability to provide an immune evasion phenotype.
4. **The PD-L1 receptor.** The PD-L1 receptor, encoded by the CD274 gene, plays a major role in suppressing the adaptive immune system. It has been reported how overexpression of PD-L1 in tumor cells leads to impaired recruitment of immune effectors^40^. We therefore considered CD274 copy number amplification as a genetic mechanism of immune evasion. We defined a CD274 copy number amplification event as samples with CD274 minimum tumor copy number greater than 3 times the average sample ploidy.
5. **The CD58 receptor.** The CD58 receptor plays an essential role in T-Cell recognition and stimulation. It has been extensively reported that CD58 alterations in B-cell lymphomas lead to immune evasion^41^. Moreover, a recent study identified CD58 loss as one the major effectors of impaired T-Cell recognition^42^. Consequently, we considered inactivation events (see above) in CD58 as alterations able to provide an immune escape phenotype.
6. **Epigenetic driven immune escape.** It has been recently reported how SETDB1 amplification leads to epigenetic silencing of tumor intrinsic immunogenicity^43^. SETDB1 amplification was recurrently found across several cancer types and therefore was considered in this study as a mechanism of immune evasion. We defined a SETDB1 copy number amplification event as samples with SETDB1 minimum tumor copy number greater than 3 times the mean sample ploidy.

A summary table with all the 21 considered genes, their associated pathway, references and their type of somatic alterations is presented in Supp. Table 1.

#### Primary and metastatic prevalence

The prevalence of a pathway alteration for a particular cohort was calculated as the number of samples with, at least, one alteration in the pathway divided by the total number of cohort samples. The presence of a genetic immune alteration in a given sample was annotated if there was, at least, one pathway with an alteration in that sample.

For the primary versus metastatic comparison, we performed a tumor-type specific Fisher’s exact test comparing pathway-specific and global escaped status prevalence across the two cohorts. P-values were adjusted with a multiple-testing correction using the Benjamini–Hochberg procedure (alpha=0.05).

### Positive selection

#### Somatic point mutations and indels

Positive selection analysis based on somatic point mutation and small indels was performed using dNdScv using the hg19 reference genome. The analysis was performed in a cohort specific, cancer type and pan-cancer manner across the two datasets. The analysis was restricted to datasets with sufficient representativeness (i.e., number of samples >=15). Global dN/dS ratios of the HLA-I (HLA-A, HLA-B and HLA-C) and the 16 non-HLA-I genes potentially targeted by mutations (i.e., excluding SETDB1 and CDC274 because their immune escape phenotype is associated with copy number gain, see Suppl. Table 1) were calculated in a pan-cancer manner using the *gene_list* attribute of the dndscv function.

#### Copy number variants

We devised a statistical test to assess positive selection in loss of heterozygosity (LOH), homozygous deletions (HD) and copy number amplifications (AMP) events. LOH, was defined as those genomic regions where the minor allele ploidy of this gene was below 0.3 and the major allele ploidy greater than 0.7. HD was defined as those regions with estimated minimum copy number below 0.5. Similarly, AMP were defined as those genomic regions with the minimum tumor copy number greater than 3 times the mean sample ploidy.

For a particular type of event overlapping with a gene, this test compares the number of observed samples bearing the alteration to the expected after whole genome randomization. More specifically, these are the followed steps:

1. Let us first denote *E* as the type of query alteration (i.e., LOH, HD or AMP), *S* as the a group of samples (usually samples from the same cancer type and same dataset) and *G_s_* as the genomic scale (i.e., nonfocal for segment lengths greater than 75% chromosome arm, focal for segments shorter than 75% of the chromosome arm and highly-focal for segments shorter than 3 Megabases).
2. For every sample *S*_i_ in {*S_1_,S_2_…S_T_*} we first gather the number and length of observed (*O_i_*) segments targeted by *E* within that *G_s_*. Only *E* events overlapping to autosomes are considered in this study. Samples that do not harbor any event of type *E* within that *G_s_* are ignored.
3. Next, for every sample *S_i_* we performed 10 independent randomizations (*R_i1_, R_i2_…R_i10_*) of the *O_i_* events, by randomly shuffling these events *E* along the autosomes. For that we used the shuffle function from pybedtools^44^ with the following parameters (genome=“hg19”, noOverlapping=True, excl=”sexual_chomosomes’’, allowBeyondChromEnd=False). In certain samples, with an extremely high segment load (*O_i_* > 10,000) or with mean ploidy of ∼1 (i.e., monoploid genome), the *noOverlapping* flag was set to False because the randomization would not converge.
4. We then binned the autosomes into 28,824 bins of 100 Kbs and counted for each bin *k_j_ {k_1_…k_28842_}* the total number of observed events *O_Tj_* as the the sum of observed events O_1k_…O_TK_ overlapping with that bin across all *S* samples.
5. Similarly, for each *R_i-th_* (R_1_…R_10_) randomization and each bin *k_j_ {k_1_…k_28842_}*, we counted the total number of simulated events as the sum of events -in that *i-th* randomization and overlapping with that bin across all samples in *S*.
6. We then performed a bin-specific comparison of the *O_Tk_* with the average number of simulated events *R_TK_* across the 10 simulations and performed a statistical test of significance using a G-test goodness-of-fit. Since chromosome starting bins were highly depleted in the simulated group (R*_TK_*), we also computed the global simulated average across all bins *k_j_ {k_1_…k_28842_}*, and used this as the expected number of events for the statistical significance assessment.
7. The p-values were adjusted with a multiple-testing correction using the Benjamini–Hochberg procedure (alpha=0.05).
8. For each gene, overlapping with one or with multiple k_j_ bins, we used the miminimal adjusted p-value significance of the bin(s) overlapping with the genomic location of the specific gene coding sequence. Therefore, by definition, two genes sharing the same bins would have similar q-value. We used ENSEMBLv88 to perform the annotation of gene exonic regions to hg19 genomic coordinates.

We observed that LINE insertions near the HLA-I locus (LINE activation site at chr6:29,920,000) in some esophageal cancer samples had an incorrect copy number estimation due to multiple insertions originating from almost the same site in the same sample. Consequently, these samples were not considered in the HLA-I homozygous deletions analysis.

The source code of the randomization test will be available upon publication in a peer-reviewed journal.

#### Distribution of mutations in HLA-I genes

LILAC mapped the HLA-A, HLA-B and HLA-C somatic mutations into the inferred HLA-I alleles (see LILAC section). LILAC provides the consequence type and coding sequence position of HLA-I alterations, which was used to display the distribution of mutations across the HLA-I coding sequence sequence. The 34 amino acids involved in peptide presentation were gathered from our neoepitope prioritization pipeline (see below). Pfam HLA-A, HLA-B and HLA-C domains were manually downloaded from the Pfam^45^ website.

We used the *geneci()* function of dNdScv to estimate the pan-cancer dN/dS ratios, which include confidence intervals, of the HLA-I genes.

### Tumor specific neoepitopes

#### Neoepitope prioritization pipeline

##### Overview

The goal of this pipeline (named as Neo) is to provide a reliable collection of neoepitopes derived from tumor specific alterations. These alterations consider point mutations (i.e., missense variants and stop loss variants), small indels (i.e., in-frame indels and frameshift) and gene fusions (in-frame and out-of-frame fusions). The neoepitope pipeline works in 2 main steps to form a comprehensive set of neopeptide and neoepitope predictions from our DNA pipeline output:

- Determination of all novel peptides (i.e., neopeptides) from all point mutations, small indels and gene fusions.
- Calculation of allele specific presentation scores using a novel binding affinity prediction algorithm.

Although we annotate with expression information from RNA (when available), the neoepitope predictions are currently based solely on mutations found in the DNA. Hence we specifically ignore RNA events such as circular RNA, RNA editing, endogenous retroviruses and alternative splicing as we are unable to determine if these are tumor specific and hence will make neoepitopes. High confidence fusions detected in RNA but not found in DNA are also currently ignored. We also acknowledge that we miss protein level events including non-canonical reading frames, post translational amino acid modifications & proteasomal peptide splicing.

##### Identification of neopeptides

We searched for potential neoepitopes for point mutations and structural variants that meet the following criteria:

1. We included somatic point mutations and indels with coding effects (i.e., missense, frameshift, in-frame or stop loss) and a SAGE filter == “PASS”.
2. We considered in-frame and out-of-frame gene fusions. See Supp. Note 2 for further details about the filtering of fusions.

Subject to the criteria above, all transcripts (or combination of transcripts in the case of fusions) are considered as candidate neoepitopes. Where 2 transcripts (or transcript combinations) lead to either the same amino acid sequence or the amino acid sequence of one transcript forms a subset of another, the transcripts are merged to form a single neoepitope. For each unique neoepitope, our pipeline outputs the amino acid (AA) sequence string broken up into ‘upstream’, ‘novel’ and ‘downstream’ segments.

Neo further annotates each of the candidate neoepitopes with TPM and direct RNA fragment support for the novel amino acid sequence. TPM per transcript is sourced from Isofox (https://github.com/hartwigmedical/hmftools/tree/master/isofox) if RNA-Seq is available or if not available is estimated as the median of the cancer type (or full cohort where cancer type is not known). Neo also reanalyses the RNA BAM to count the RNA depth at the location of the variant that caused the neoepitope and the direct RNA fragment support for the neoepitope (defined as matching precisely the 1st novel AA and 5 bases either side). See Supp. Note 2 for further information about identification candidate neoepitopes from the tumor analytical pipeline.

##### Calculation of allele specific binding affinity and presentation scores

Using the identified neoepitopes, we determine all candidate peptide and allele pairs (pHLA) combinations that may be presented by the cell. For each candidate neoepitope, we consider all peptides between 8 and 12 length which either overlap the novel amino acid sequence or overlap both the upstream and downstream amino acid sequence.

For each pHLA we estimated a presentation likelihood based on a newly developed Position Weighted Matrix (PWM) algorithm that considers both the binding affinity and the processing likelihood for each pHLA pair. Supp. Note 2 describes in detail all the steps and validation of our newly developed pipeline.

Each pHLA score was then ranked compared to a random set of 100,000 peptides derived from the human canonical proteome to derive a presentation likelihood rank for each pHLA.

##### Expression adjusted presentation likelihood algorithm

Multiple studies have shown the importance of including RNA expression in the prioritization of neoepitopes. The pHLA presentation scores were further adjusted by the inclusion of the normalized RNA expression of the mutated transcript/s (including the 5’ and 3’ transcripts in gene fusions). When RNA expression was not available, we used the average expression across the patient cancer type or the pan-cancer when the cancer type is unknown. For further details about the logic, validation and how we estimated the RNA expression of each pHLA see Supp. Note 2. We then calculated the expression-adjusted likelihood rank (ExpLikelihoodRank) for each pHLA using the expression adjusted likelihood compared to the same randomly selected peptides.

Finally, we defined the collection of neoepitopes as those pHLA with a LikelihoodRank < 0.02 and ExpLikelihoodRank < 0.02 (i.e., within the 2% percentile of all peptides). Hence, the total neoepitope load of a patient tumor sample is the sum of all pHLA neoepitopes with LikelihoodRank < 0.02 and ExpLikelihoodRank < 0.02.

#### Neoepitope clonality

Each predicted neoepitope (see above) derived from point mutations and small indels was matched with the source variant estimated clonality from PURPLE. We defined clonal mutations as those with a subclonal score lower than 0.85. Gene fusions were not considered in this analysis.

#### Calculation and randomization of neoepitope ratio

We wanted to evaluate whether LOH HLA-I tends to select the HLA-I allele with highest neoepitope repertoire. Let us first introduce the neoepitope allele ratio (*nr*). Given an HLA-I gene, G, we defined *nr* as: 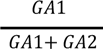, where G_A1/2_ is the number of predicted neoepitopes of allele 1 and allele 2, respectively. For each patient tumor sample, the assignment of allele number (i.e., allele 1 or allele 2) was randomly performed. Next we followed the next steps:

1. For each patient sample with LOH HLA-I we calculated the *nr* across the HLA-I genes targeted by the LOH. Homozygous HLA cases were not considered, as they *nr* is by definition 0.5.
2. We then grouped the *nr* into 8 buckets: [0.0-0.35), [0.35, 0.4), [0.4, 0.45), [0.45, 0.5). [0.5, 0.55), [0.55, 0.6), [0.6, 0.65) and [0.65, 1.0]. Consequently, each bucket included N allele pairs with a *nr* within the bound limits.

1. Next, we performed 100 bootstraps by randomly subsampling 75% of the total number of available allele pairs.
2. For each bootstrap iteration *i-th* (*i* ∈ 1..100) and each bucket we estimated the frequency of allele 1 loss (F_A1_loss_) as the number of cases with allele 1 loss compared to the total number of cases in that bucket. Similarly, we computed the expected frequency (F_A1exp_) by randomly assigning LOH events to the allele1.
3. We then computed the bucket-specific average and standard deviation of F_A1loss_ and F_A1exp_ values across the 100 bootstraps.
4. Finally, we performed a Kolmogorov–Smirnov test to compare the observed distribution to the expected given random distribution of LOH events.

This test was applied to LOH HLA-I, focal LOH HLA-I and non-focal LOH HLA-I events across the metastatic (Hartwig) and primary (PCAWG) datasets.

### Tumor genomic features and GIE risk

We aimed at identifying cancer type genomics features associated with an increased GIE risk. Hence, we performed a cancer type aggregation of the two datasets (i.e., metastatic and primary) to increase statistical power of the analysis. We then computed 95 genomic features and evaluated its association with GIE across 32 cancer types with sufficient representativeness (i.e., total number of samples >=15).

#### Tumor mutation burden, neoepitope load and SV load

For each cancer type, we used a univariate logistic regression to quantify the association of 21 TMB-related measurements (see Supp. Table 6 for full list of evaluated features) and the presence/absence of a GIE event. Independent variables were z-scored. The *Logit()* function (with default parameters) from the statsmodels v.0.13.1 library was used to perform the logistic regression. This function provides the odds ratio with confidence intervals alongside the p-value of significance. The p-values were adjusted with a multiple-testing correction using the Benjamini–Hochberg procedure (alpha=0.05).

The clonality of each variant was defined using the PURPLE subclonal likelihood estimation. More specifically, a variant was considered as clonal if the estimated subclonal likelihood was lower than 0.85.

The global neoepitope load of each patient’s sample was calculated as the sum of the predicted neoepitopes (i.e., allele specific neoepitope repertoire, see methods Tumor specific neoepitopes) across the germline HLA-A alleles inferred by LILAC. The subset of neoepitopes (i.e., fusion derived, mutation derived, clonal and subclonal) was computed by matching the source alteration -and their clonality-of each predicted neoepitope. Therefore, a mutation (or gene fusion) may be the source for multiple neoepitopes.

#### Mutational signatures

The number of somatic mutations falling into the 96 single nucleotide substitution (SBS), 78 double base substitutions (DBS) and 83 indel (ID) contexts (as described in the COSMIC catalog^46^ https://cancer.sanger.ac.uk/signatures/) was determined using the R package mutSigExtractor (https://github.com/UMCUGenetics/mutSigExtractor, v1.23).

SigProfilerExtractor (v1.1.1) was then used (with default settings) to extract up to 21, 8 and 10 de novo mutational signatures for SBS, DBS and indels (respectively). This was performed separately for each of the 22 tissue types which had at least 30 patients in the entire dataset (aggregating primary and metastatic samples, see Supp. Table 3). Tissue types with less than 30 patients as well as metastatic patients with unknown primary location type were combined into an additional ‘Other’ group, resulting in a total of 23 tissue types for signature extraction. In order to select the optimum rank (i.e. the eventual number of signatures) for each tissue type and mutation type, we manually inspected the average stability and mean sample cosine similarity plots output by SigProfilerExtractor. As a result, there were 484 *de novo* signature profiles extracted across the 23 tissue type groups (see Supp. Table 3 and Supp. Table 6). Least squares fitting was then performed (using the fitToSignatures() function from mutSigExtractor) to determine the per-sample contributions to each tissue type specific *de novo* signature.

The extracted *de novo* mutational signatures with high cosine similarity (>=0.85) to any reference COSMIC mutational signatures with known cancer type associations^46^ were labeled accordingly (number of labeled *de novo* signatures = 271 matched to 57 COSMIC references).

For the collection of remaining non-labeled *de novo* signature profiles (number of unlabeled *de novo* signatures =213) of each mutation type, we reasoned that there could be one or more signatures that are highly similar to those found in the set of signatures of other tissue types (and thus likely representing the same underlying mutational process) and that have not been yet matched to a COSMIC reference. We therefore performed clustering to group likely equivalent signatures and to label them as such. Specifically, we followed the next steps:

1. We calculated the pairwise cosine distance between each of the *de novo* signature mutational profiles.
2. We performed hierarchical clustering and used the base R function cutree() to group signature profiles over the range of all possible cluster sizes (min no. clusters = 2; max no. of clusters = number of signature profiles for the respective mutation type).
3. We next calculated the silhouette score at each cluster size to determine the optimum number of clusters.
4. Finally, we grouped the signature profiles according to the optimum number of clusters. This yielded in total 47 *de novo* signature clusters (see Supp. Table 6).

For certain *de novo* signature clusters we were able to manually assign the potential etiology by relying on the average similarity to the COSMIC reference mutational signatures. For instance, SBS_denovo_clust_1 represented a collection of *de novo* signatures highly similar to the reference SBS2 and SBS13 from COSMIC, linked to APOBEC mutagenesis. In many cases the mutational signatures displayed an aggregation of both mutational spectra (SBS+SBS13) preventing the reference annotation in the first step of our pipeline. Similarly, DBS_denovo_clust_5 represented a collection of *de novo* signatures similar to the DBS5 of COSMIC, which had been linked to platinum treatment exposure. These *de novo* mutational signatures presented the characteristic CT>[AA or AC] peak of DBS5 COSMIC signature in combination with residual contribution from other DBS channels. Finally, we assigned MMR deficiency as the etiology for several clusters (e.g.., ID_denovo_clust_1, see Supp. Table 6) as these clusters were enriched in MMR deficient samples.

Next, for each cancer type, we used a logistic regression to quantify the association between the number of somatic point mutations, indels or double base substitutions (DBS) attributed to a certain mutational signature and the presence/absence of a GIE event. We only considered mutational signatures with known/suspected etiology or with high similarity to a reference COSMIC signature (cosine similarity >=0.85) as well as high incidence in a particular cancer type (i.e., at least 20 samples with a mutational signature exposure greater or equal than 100 mutations). 49 mutational signatures fulfilled these filters in at least one cancer type. Moreover, to diminish associations that could be mainly attributed to an elevated molecular age, we also included the exposure to aeing mutational signature(s) as an independent variable (SBS1 + SBS5 cumulative exposure as a proxy for molecular age). Independent variables were z-scored. The *Logit()* function (with default parameters) from the statsmodels library was used to perform the logistic regression. This function provides the odds ratio with confidence intervals alongside the p-value of significance for both dependent variables. The p-values were adjusted with a multiple-testing correction using the Benjamini–Hochberg procedure (alpha=0.05).

#### MMR and HR deficiency

We also tested whether Mismatch repair deficiency (MMRd) and Homologous repair deficiency (HRd) were predictive of GIE. The Hartwig analytical pipeline provides the MMRd status of each processed tumor sample (i.e., microsatellite stable or microsatellite unstable). Analogously, the CHORD^47^ software was used to evaluate HRd in tumor samples. Fisher’s exact test was used to evaluate the significance. A minimum of 5 DNA repair-deficient tumor samples were required to assess the significance. P-values were adjusted with a multiple-testing correction using the Benjamini–Hochberg procedure (alpha=0.05).

#### DNA viral insertion and Whole Genome duplication

The presence of viral DNA and whole genome duplication (WGD) is provided by the Hartwig analytical pipeline. A Fisher’s exact test was used to evaluate the significance. A minimum of 10 tumor samples harboring viral DNA insertions were required to assess the significance. P-values were adjusted with a multiple-testing correction using the Benjamini–Hochberg procedure (alpha=0.05).

#### Immune infiltration deconvolution

For samples with available tumor RNA-Seq data we performed an immune infiltration deconvolution based on the normalized TPM and RPKM values in Hartwig and PCAWG, respectively. More specifically, we implemented 6 different markers of immune infiltration: the natural killer cells (NK) quantification by Patrick Danaher et al.^48^, the global immune infiltration, CD8^+^ T-cells and CD4^+^ T-cells implemented by Teresa Davoli et al.^38^, the T-cell infiltration used by Catherine Grasso et al.^4^ and the preliminary IFN-γ profile reported by Mark Ayers et al^49^.

Next, we used univariate logistic regression to quantify the association of these measurements with GIE prevalence. Independent variables were z-scored. The *Logit()* function (with default parameters) from the statsmodels library was used to perform the logistic regression. This function provides the odds ratio with confidence intervals alongside the p-value of significance for both dependent variables. The p-values were adjusted with a multiple-testing correction using the Benjamini–Hochberg procedure (alpha=0.05).

#### HLA-I supertypes

We performed a cancer-type specific Fisher’s exact test to assess enrichment of HLA-I supertypes with the GIE frequency. Only HLA-I supertypes present in at least 50 patients were evaluated. HLA-I supertypes were gathered from ref.^50^ and manually curated. P-values were adjusted with a multiple-testing correction using the Benjamini–Hochberg procedure (alpha=0.05).

#### HLA-I divergence

We calculate the germline average and cumulative HLA-I divergence^51^ as the mean and sum of LILAC’s HLA-I alleles pairwise divergence. Both measurements were independently regressed against the GIE prevalence in a cancer type specific manner. Following the same methodology used with other features, a logistic regression was used to evaluate the significance of the association.

#### Pre-biopsy treatment exposure

We also tested whether exposure to pre-biopsy treatment had a predictive value for GIE prevalence. For this analysis, we only relied on metastatic pre-treated samples with available pre-treatment information (N=2,272). Two treatment groups were tested: chemotherapy and immunotherapy because of the commonness across cancer types. A minimum of 20 treated samples were required to carry out the association between a treatment and GIE in a particular tumor type. Fisher’s exact test was used to evaluate the significance. P-values were adjusted with a multiple-testing correction using the Benjamini–Hochberg procedure (alpha=0.05).

#### GIE and TMB dependence

We aggregated the two datasets, metastatic and primary, to increase the robustness of this analysis. We then defined 20 evenly arranged buckets of the log_10_(TMB) scale, starting from the 1^th^ percentile and ending in the 99^th^ percentile values. Next, each sample with a log_10_(TMB) equal to S_tmb_, was allocated to the *i-th* (*i* ∈ 1..20) bucket such as log_10_(TMB)_i-1_ < S_tmb_ <= log_10_(TMB)_i_. Samples with a S_tmb_ greater than the last bucket threshold (i.e., log_10_(TMB)_20_) were allocated into the last bucket. The number of mutations of each bucket was displayed as the number of mutations per megabase (Muts/Mb) by dividing the total number of mutations by 3,000 (i.e., approximated number of human genome megabases). Finally, the GIE frequency (GIE_freq_) of the *i-th* bucket was defined as the number of GIE samples in the *i-th* bucket compared to the total number of available samples in that bucket.

In order to enable the calculation of the uniformity in GIE frequency among samples in the sample bucket, we performed N (N=1,000) bootstraps of the 50% of samples allocated to each bucket. We then calculated the average and standard deviation of the GIE_freq_ across the bootstraps.

A similar approach was conducted to analyze the relationship between the predicted neoepitope load and GIE frequency. The number of neoantigens of each bucket was estimated as the 1%^32^ and the 5%^6^ of the total predicted neoepitopes assigned to that bucket threshold.

## Data availability

The Hartwig dataset used in this study are freely available for academic use from the Hartwig Medical Foundation through standardized procedures and request forms that can be found at https://www.hartwigmedicalfoundation.nl/en/applying-for-data/. This includes raw sequencing data (BAM files and unmapped reads) as well as the processed data through the latest version of the Hartwig tumor processing pipeline.

To access to the PCAWG dataset processed by the Hartwig Medical Foundation analytical pipeline researchers will need to apply to the TCGA data access committee via dbGaP (https://dbgap.ncbi.nlm.nih.gov/aa/wga.cgi?page=login) for the TCGA portion of the dataset, and to the ICGC data access compliance office (http://icgc.org/daco) for the ICGC portion of the dataset.

Raw sequencing data of the high-resolution HLA typing performed by GenDx will be made available upon acceptance in a peer-reviewed journal.

## Code availability

The Hartwig analytical processing pipeline is available at (https://github.com/hartwigmedical/pipeline5) and implemented in Platinum (https://github.com/hartwigmedical/platinum). LILAC’s source code is available at (https://github.com/hartwigmedical/hmftools/tree/master/lilac). The source code of the neoepitope prioritization pipeline and the code used to prepare the figures will be made public upon acceptance in a peer-reviewed journal.

## Acknowledgements

This publication and the underlying study have been made possible partly on the basis of the data that Hartwig Medical Foundation and the Center of Personalised Cancer Treatment (CPCT) have made available to the study. We thank Dr. Arne Van Hoeck and Dr. Abel Gonzalez Perez for critical reading of the manuscript and their valuable input. We also thank Dr. Arne Van Hoeck for the assistance in the dataset collection and Luan Nyguen for the help in the mutational signature extraction. Finally, we appreciate the cooperation of Wietse Mulde, Maarten Penning and Hanneke Merkens in LILAC’s orthogonal validation. Biorender templates were retrieved from https://app.biorender.com/biorender-templates.

## Author contributions

Conceptualization, F.M.J, E.C and P.P. Methodology, F.M.J, P.P and C.S. Software, P.P. and C.S. Validation F.M.J., P.P and E.R. Formal Analysis, F.M.J. and P.P. Investigation F.M.J. and P.P. Resources E.C. and P.P. Data Curation, F.M.J, P.P, C.S and E.R. Writing Original Draft, F.M.J. Writing - Review & Editing E.C. and P.P. Visualization F.M.J. Supervision F.M.J and E.C. Project Administration, F.M.J. and E.C. Funding Acquisition E.C.

## Correspondence

Correspondence should be addressed to Francisco Martínez-Jiménez and Edwin Cuppen.

## Supplementary Information

### Supplementary Figure legends

**Supplementary Figure 1.**
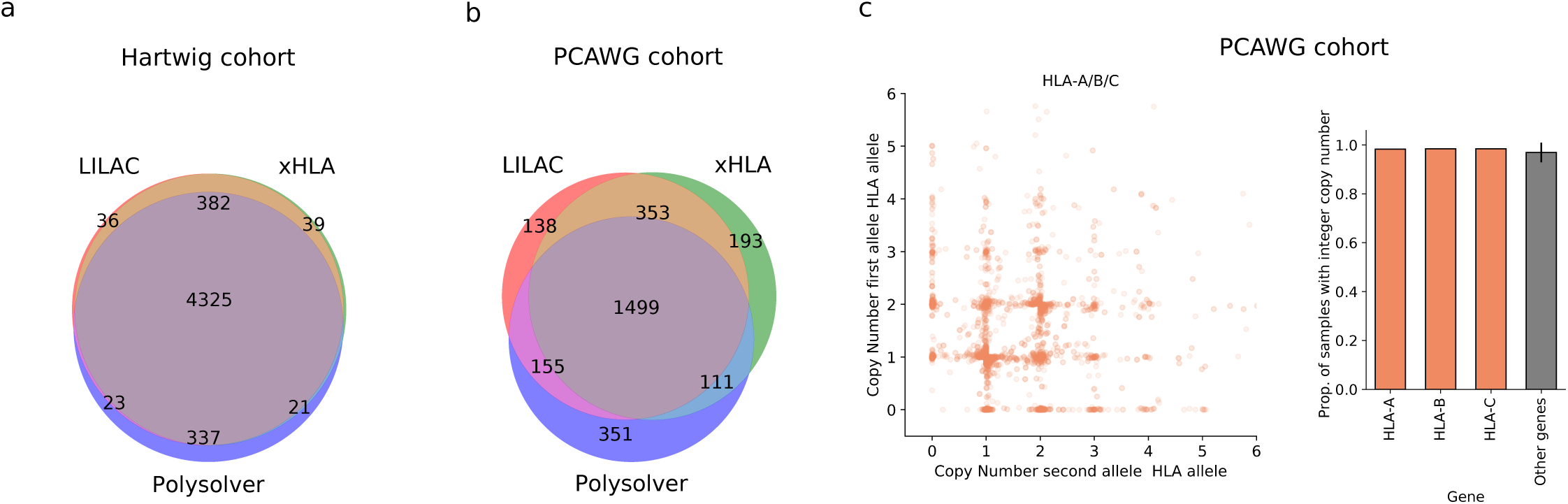
LILAC validación. Venn diagrams representing the overlap of germline samples with perfect 2-field HLA-I allotype match in a) Hartwig and b) PCAWG. c) Left, copy number of minor and major alleles of HLA-A, HLA-B and HLA-C in PCAWG. Right, proportion of samples with integer copy number of HLA-I genes (orange) compared to the average of 1,000 random genes in PCAWG (gray). The error bar represents the standard deviation.

**Supplementary Figure 2.**
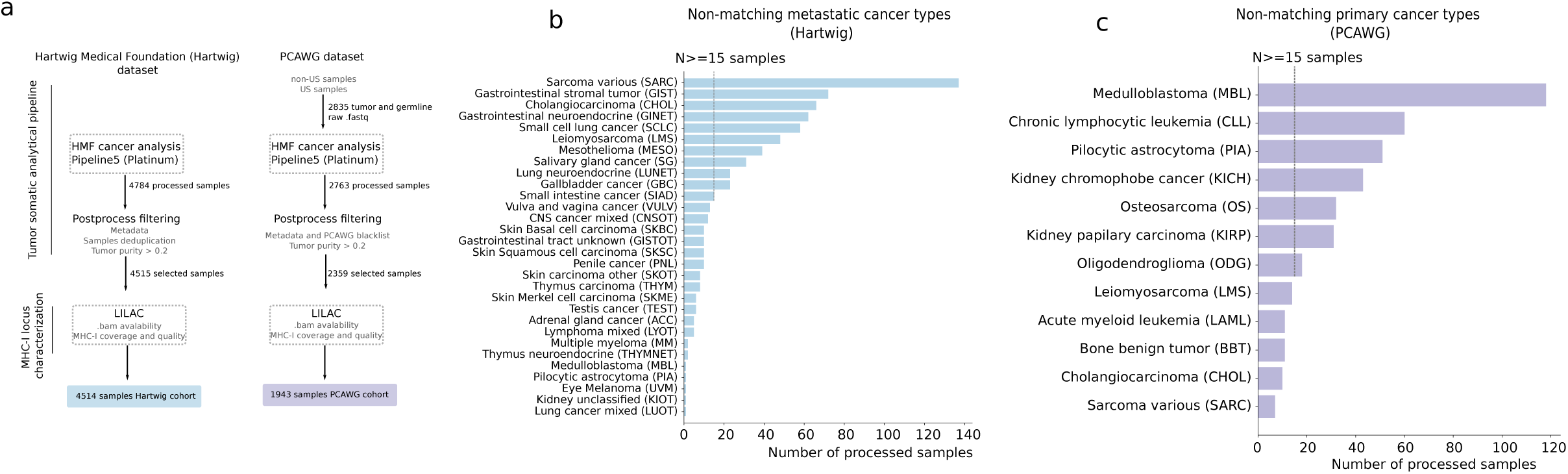
GIE in primary and metastatic tumors. a) Workflow of the processing pipeline used in this study in Hartwig (left) and PCAWG (right). Each rectangle represents a processing step. The resulting number of selected samples are displayed at the bottom. b) number of metastatic (Hartwig) samples across cancer types that lack sufficient representation in the primary (PCAWG) dataset. c) analogous representation for primary (PCAWG) samples. Vertical dashed lines (N>= 15 samples) represent the threshold of samples to consider a cohort as sufficiently populated.

**Supplementary Figure 3.**
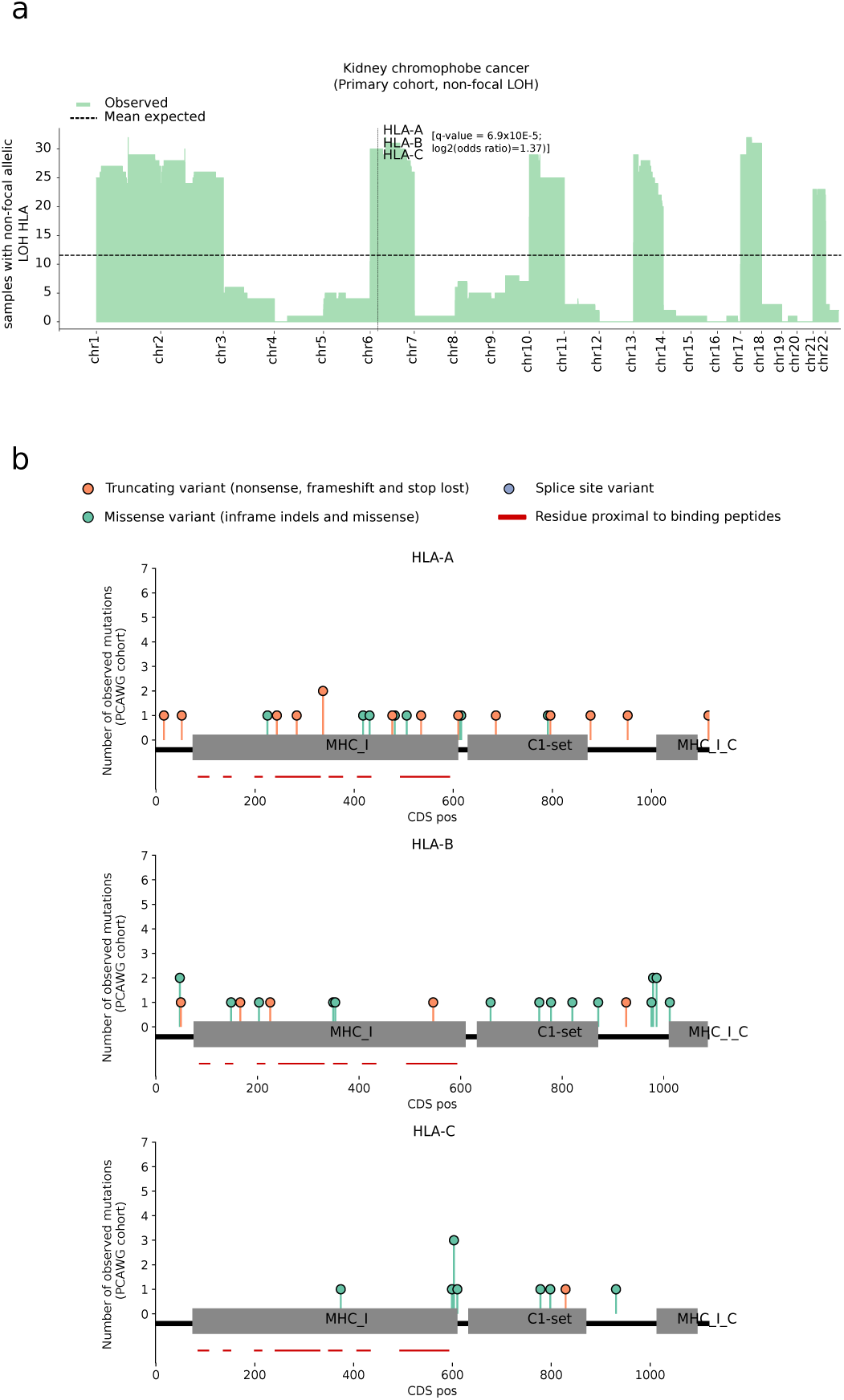
Positive selection of HLA-I. a) Distribution of non-focal LOH events along the autosomes in the primary kidney chromophobe cancer cohort. X-ticks represent the chromosomal starting position. Dashed horizontal lines represent the expected mean after randomization. Vertical dashed lines highlight the HLA-I genomic location. b) Needle plots representing the pan-cancer distribution of somatic mutations along the HLA-A, HLA-B and HLA-C protein sequences in the primary dataset. Mutations are coloured according to the consequence type. CDS pos, coding sequence position.

**Supplementary Figure 4.**
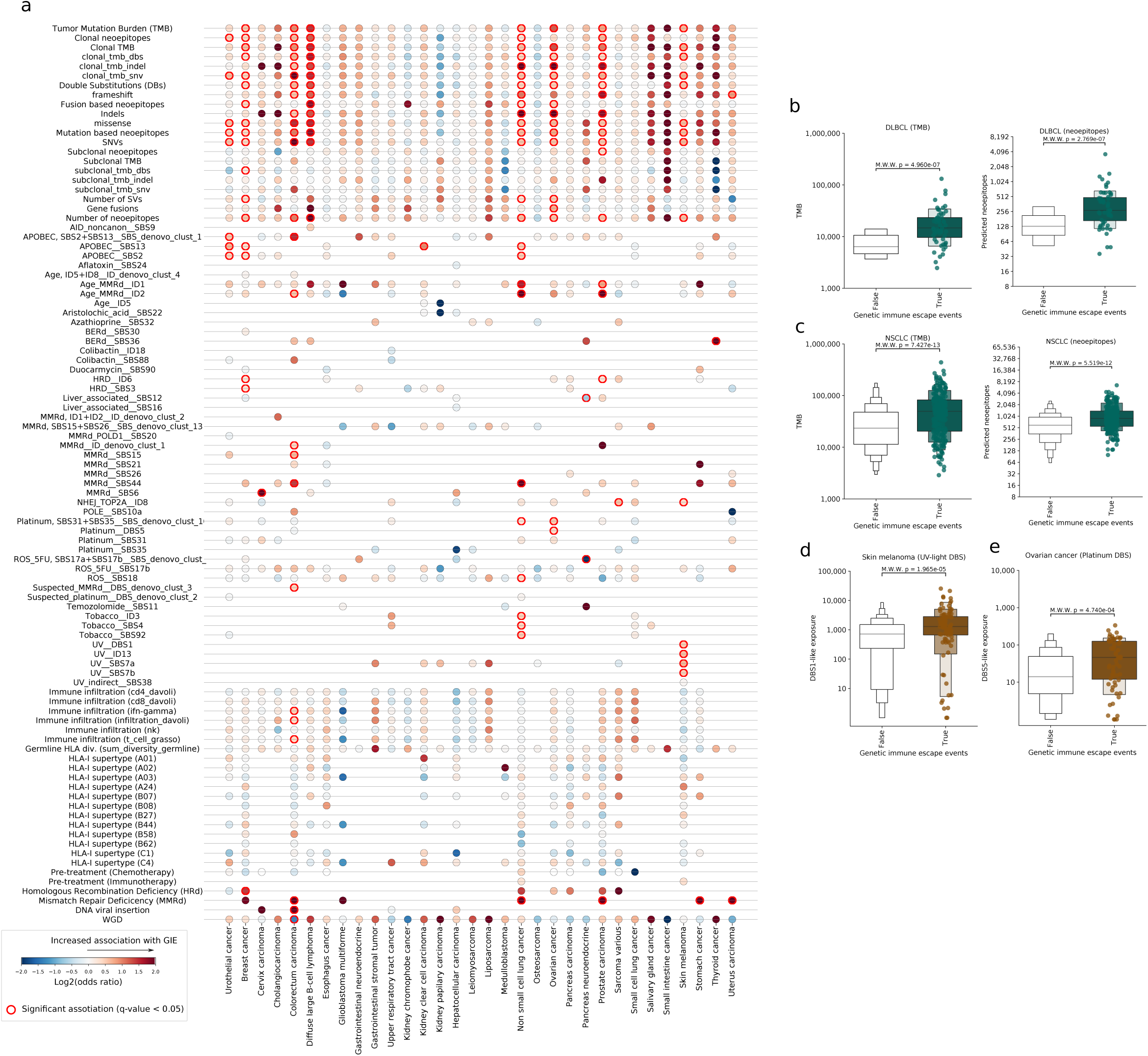
GIE association with cancer genomic features. a) Heatmap displaying the association of 95 genomic features with GIE frequency across 32 cancer types. Significant associations are highlighted by a red border line. Dot colors are coloured according to the Log_2_(odds ratio). b) Comparison of the TMB (left) and predicted neoepitope load (right) between samples bearing GIE alterations in DLBCL and c) NSCLC. d) Comparison of the UV-light attributed DBS between samples bearing GIE alterations and wild-type (non-GIE) in skin melanoma samples. e) Similar comparison for DBS attributed to platinum treatment in ovarian cancer. Box-plots: center line, median; box limits, first and third quartiles; whiskers, lowest/highest data points at first quartile minus/plus 1.5× IQR. M.W.W, Mann–Whitney U test p-value. DBS, double base substitutions.

**Supplementary Figure 5.**
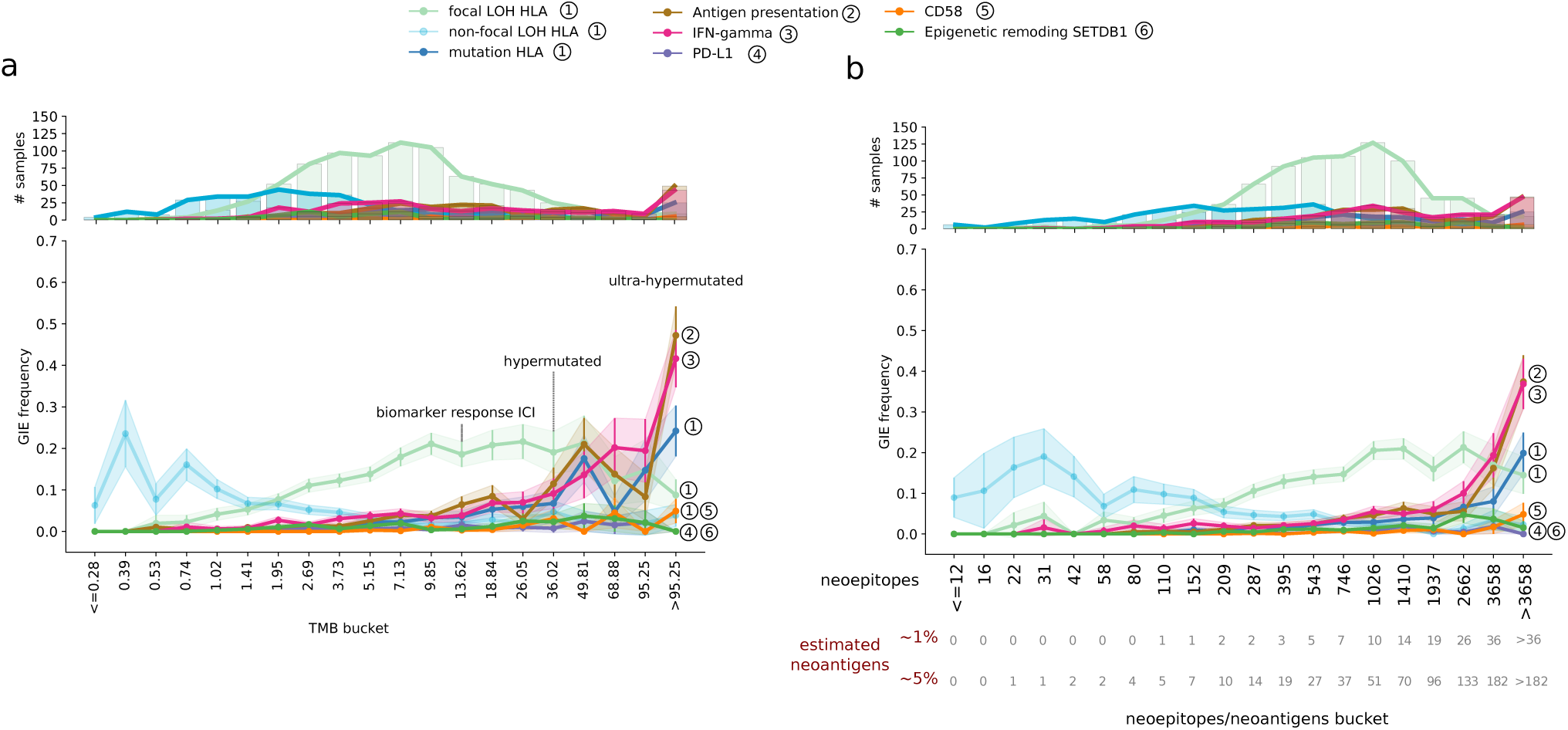
Immune evasion mechanism and TMB. a) Top panel, bar plots representing the number of mutated samples per TMB bucket split by GIE pathway and mechanism. Bottom panel, representation of pathway-specific GIE frequency across twenty evenly distributed TMB buckets. Dots represent the average GIE frequency across 1,000 bootstraps. Error bars and the shades represent ± standard deviation. The annotated numbers are associated with the identifier of the immune escape pathway, relative to Figure 1a. b) Using predicted neoepitopes as baseline buckets. Bottom labels, number of estimated neoantigens as a relative percentage (1% and 5%) of the number of predicted neoepitopes in the bucket. Muts/Mb, mutations per megabase.

### Supplementary Tables

**Supplementary Table 1. Pathways and genes involved in genetic immune escape.**

**Supplementary Table 2. HLA-I typing of Hartwig and PCAWG patients and orthogonal validation of LILAC.**

**Supplementary Table 3. Datasets metadata and cancer type representation.**

**Supplementary Table 4. Sample specific GIE annotation and cohort-wise GIE frequency.**

**Supplementary Table 5. Positive selection in HLA-I and non HLA-I genes.**

**Supplementary Table 6. Genomics features and GIE association.**

### Supplementary Notes

**Supplementary Note 1. LILAC.**

**Supplementary Note 2. Neoepitope prioritization pipeline (Neo).**

