## Supplementary Note 1 for "Genetic immune escape landscape in primary and metastatic cancer"

#### Lilac

LILAC is a WGS tool to determine the HLA Class I types for the germline of each patient as well as determining the status of each of those alleles in the tumor including complete loss of one or more alleles, allele specific somatic mutations and allelic imbalance.

LILAC uses the IMGT/HLA[1] database (https://www.ebi.ac.uk/ipd/imgt/hla/download.html) as a reference set of Human MHC class I alleles, and performs typing to 4-digits, which means it uniquely identifies a specific protein, but ignores synonymous variants (6 digits) and intronic differences (8 digits).

There are many existing tools which can perform HLA class 1 typing notably Polysolver[2], xHLA[3] and Optitype[4]. LILAC offers a number of potential advantages including:

- Estimated 99.8% accuracy on 30x - 100x WGS, with improved accuracy particularly for rare alleles.
- Fully integrated analysis of paired tumor-normal sample data to call allele-specific copy number and assignment of somatic variants to specific alleles.
- Detection of novel germline variants / alleles (including indels) via analysis of unmatched fragments
- Identification of HLA-Y presence (a pseudogene with high similarity to HLA-A which is present in up to 20% of the population but is not present in the reference genome)

LILAC is designed to take SAGE (https://github.com/hartwigmedical/hmftools/blob/master/sage/README.md) somatic point mutation output and PURPLE (https://github.com/hartwigmedical/hmftools/blob/master/purple/README.md) predictions of allele specific copy numbers as input. These can also be supplied by other point mutation or copy number callers.

LILAC supports GRCH37, hg19 and hg38 (no alt) ref genomes. Note that if the genome has been aligned to an hg38 genome with HLA alt contigs, that fragments from these contigs will need to be extracted and realigned to the no alt genome prior to running LILAC.

LILAC has been optimized to 30-40x germline coverage and 100x tumor coverage with paired 151 base reads. Shorter reads or lower depth coverage may be problematic. LILAC will achieve accurate results on tumor samples with sufficient coverage (ideally >=100x), but in tumors with high purity and LOH, the lost allele in the tumor may be difficult to detect.

### Configuration

To install, download the latest compiled jar file from the download links (https://github.com/hartwigmedical/tools/tree/master/lilac#version-history-and-download-links)

### Resource Files

Lilac requires the following resource files:

- Ref Genome .fasta
- HLA allele nucleotide and amino acid definitions - available from the Hartwig resources page, or can be generated by Lilac (see below). The current version used by Lilac is IPD-IMGT/HLA 3.42.0.
- Allele Frequencies - taken from Hartwig's cohort of ~5000 samples, available from the Hartwig resources page

Reference files are available here HMFTools-Resources (https://resources.hartwigmedicalfoundation.nl/): then navigate to 'Lilac'

##### Generate HLA Definitions

Nucleotide and amino acid allele definitions are available here:

- https://www.ebi.ac.uk/ipd/imgt/hla/download.html
- ftp://ftp.ebi.ac.uk/pub/databases/ipd/imgt/hla/

The required files are A_nuc.txt, B_nuc.txt, C_nuc.txt, Y_nuc.txt, A_prot.txt, B_prot.txt and C_prot.txt.

Convert these definition files to the reference files used by Lilac with the following command:

```
java -cp lilac.jar com.hartwig.hmftools.lilac.utils.GenerateReferenceSequences \
   -resource_dir /path_to_definition_files/ \
   -output_dir /output_dir/ \
```

#### Mandatory Arguments

| Argument | Description |
|---|---|
| sample | Sample ID |
| ref_genome | Reference genome fasta file |
| reference_bam | Sample's germline BAM |
| resource_dir | Path to Lilac resource files, ie hla_ref_aminoacid_sequences.csv, hla_ref_nucleotide_sequences.csv and lilac_allele_frequencies.csv. |

NOTE: Lilac handles BAMs which have been sliced for the HLA gene regions.

If a sample's tumor BAM is provided in place of the reference BAM, then Lilac will determine the allele solution from it instead.

#### Optional Inputs

| Argument | Description |
|---|---|
| ref_genome_version | V37 (default), V38 or HG19 (ie 37 with 'chr' prefix) |
| tumor_bam | Sample's tumor BAM |
| rna_bam | Sample's RNA BAM if available |
| gene_copy_number_file | Sample gene copy number file from Purple |
| somatic_variants_file | Sample's somatic variant VCF file, for annotation of HLA gene variants |

#### Optional Arguments

| Argument | Default | Description |
|---|---|---|
| min_base_qual | 30 | Min base quality for BAM reads, see documentation for details |
| min_evidence | 2 | Min absolute required fragments for evidence phase |
| min_evidence_factor | 0.00075 | Min relative required fragments for evidence phase, as a fraction of total fragments |
| min_high_qual_evidence_factor | 0.000375 | Minimum relative required high base-quality fragments for evidence phase, as a fraction of total fragments |
| min_fragments_per_allele | 7 | See documentation for details |
| min_fragments_to_remove_single | 40 | See documentation for details |
| top_score_threshold | 5 | Maximum difference in candidate solution score vs top score as a percentage of total fragments |
| log_debug | Off (logs at INFO) | Logs in verbose mode |
| debug_phasing | Off | Logs phasing evidence construction |

| | | |
|---|---|---|
| expected_alleles | Not applied | List of alleles separated by ';'. These alleles will have their coverage and ranking reported even if not in the winning solution |
| restricted_alleles | Not applied | List of alleles separated by ';'. Restrict evaluation to only these alleles. |
| threads | 1 | Number of threads to use for complex evaluation |

#### Example Usage

```
java -jar lilac.jar \
   -sample COLO829T
   -ref_genome /path_to_ref_genome_fasta_file/ \
   -resource_dir /path_to_lilac_resource_files/ \
   -reference_bam /sample_data_path/COLO829R.bam \
   -output_dir /output_dir/ \
```

Or with tumor BAM, somatic variants VCF and Purple gene copy number inputs:

```
java -jar lilac.jar \
   -sample COLO829T
   -ref_genome /path_to_ref_genome_fasta_file/ \
   -resource_dir /path_to_lilac_resource_files/ \
   -reference_bam /sample_data_path/COLO829R.bam \
   -tumor_bam /sample_data_path/COLO829T.bam \
   -somatic_vcf /sample_data_path/COLO829T.purple.somatic.vcf.gz \
   -gene_copy_number /sample_data_path/COLO829T.purple.cnv.gene.tsv \
   -output_dir /output_dir/ \
```

#### Algorithm

The starting point for the LILAC algorithm is the complete set of possible 4 digit alleles and all the fragments aligned to HLA-A, HLA-B and HLA-C. Where multiple 6 digit or 8 digit types are present in the IMGT/HLA database, LILAC uses the numerically lowest type for all calculations. Note that 2 HLA-A alleles {A*31:135, A*33:191} have been removed from the database due to it frequently being found as a low level artifact (likely due to the high similarity to closely related genes and pseudogenes such as HLA-H).

LILAC algorithm begins with collecting all fragments which are not duplicates and have:

- At least one read with an alignment overlapping a coding base of HLA-A, HLA-B or HLA-C; and
- all alignments within 1000 bases of a HLA coding region; and
- a mapping quality of at least 1

The algorithm then has 2 main phases to determine the germline alleles: an ‘elimination’ phase which aims to remove allele candidates that are clearly not present and an ‘evidence’ phase where LILAC considers all possible sets of 6 alleles amongst the remaining candidates and chooses the solution that best explains the fragments observed.

After the germline alleles are determined, LILAC determines the tumor copy number and any somatic mutations in each allele. Note that if more than 300 bases of the HLA-A,HLA-B and HLA-C coding regions have less than 10 coverage, then LILAC will fail with errors and will not try to fit the sample.

#### Elimination phase

The elimination phase is primarily an optimization. It’s goal is simply to reduce the number of possible alleles from ~14k present in the IMGT/HLA database to a manageable number such that the evidence phase can run efficiently. The principle in the elimination phase is to remove any allele that does not have at least a certain minimal coverage of each of it’s amino acids and bases. To mitigate the chance of inadvertently eliminating a true allele, common alleles may be recovered at the end of the elimination phase if they have sufficient unique support, but are then penalized in the subsequent evidence phase relative to other candidate alleles.

The steps in the elimination phase are:

##### 1. Nucleotide matrix

At each coding position, create a matrix of (high quality) nucleotide count and determine all bases which are heterozygous across all 6 alleles.

We do this 3 separate times for HLA-A, B and C. Each time we consider fragments from any of the alignment records that have similar exon boundaries to the type in question. For instance, fragments from the earlier exons which have identical boundaries across all 3 genes will be used to construct the A, B and C matrices, but fragments from the later exons may only contribute to A and B or perhaps only C.

The counts of supporting fragments are then aggregated at each position to construct the nucleotide matrix.

Fragments with in-frame indels are only included if the indel matches an existing hla type allowing for realignment. Fragments with out-of-frame indels are always excluded (note for the special case of C*04:09N, a relatively common allele with out of frame indel, it is explicitly rescued at a later stage

During the elimination phase, nucleotide candidates are filtered to include only those with at least max(1,0.000375 * FragmentCount) high quality (base qual > min(30,medianBaseQuality)) fragment and at least max(2,0.00075 * FragmentCount) fragments overall support. Subsequently, base quality is not considered. Sites with more than 1 nucleotide candidate are deemed heterozygous and sites with only 1 are considered to be homozygous across all 6 alleles.

Any alleles with bases that do not match both the heterozygous and homozygous locations of the nucleotide matrix are eliminated.

##### 2. Amino acid matrix

Similarly to the nucleotide matrix, LILAC also constructs a matrix of amino acid candidates. Again, amino acid candidates are filtered to those where at least max(1,0.000375 * FragmentCount) fragments support with high base quality (all 3 nucleotides) and at least max(2,0.00075 * FragmentCount) fragments over all. The codon matrix can include inframe insertions and deletions where these match at least one known allele (base quality is not considered).

Exon boundary 'enrichment' is applied for all shared amino acids across all 3 genes (amino acids index < 298). This enriches any fragment with nucleotides on one side of an exon boundary with any homozygous nucleotides from the other side so that an amino acid is able to be constructed.

Similarly to the nucleotide matrix, any alleles with an amino acid or inframe indel that do not match the amino acid matrix are eliminated.

##### 3. Phased haplotypes

In this step we phase the heterozygous amino acid locations and eliminate any alleles that are not supported by phased locations with sufficient overall shared coverage.

First, we find phased evidence of each consecutive pair of heterozygous codons and record the haplotypes of all fragments containing both codons. This is performed separately for each of HLA-A, HLA-B and HLA-C to account for differences in exon boundaries. Fragments which overlap amino acid 338 onwards will only be assignable to a subset of the alleles, since the exon boundaries differ after this amino acid. The points are only phased if the total coverage is at least 7 fragments per allele [minFragmentsPerAllele] included in the subset at that location (ie. between 14 and 42 fragments with shared coverage depending on the amino acid location). A phased haplotype with only 1 supporting fragment will be removed if the total fragments supporting the pair is 40 or more [minFragmentsToRemoveSingle] (assumed to be a sequencing error).

We then iteratively choose the phased haplotype with the most support and perform the following routine:

- Find other phased evidence that overlaps it.
- Find the minimum number of codon locations required to uniquely identify each phased evidence, eg, if the left evidence has haplotypes [SP, ST] you only need the last codon but if the right evidence has haplotypes [PD, TD, TS] you would need both codon locations.
- Find evidence of fragments that contain all the required codons. As with the paired evidence, there must be at least at least 7 fragments per allele supporting the pair [minFragmentsPerAllele] and a haplotype with only 1 supporting fragment will be removed if the total fragments supporting the pair is 40 or more [minFragmentsToRemoveSingle].
- Check that the new overlapping evidence is consistent with the existing evidence.
- Merge the new evidence with the existing paired evidence.
- Replace the two pieces of used evidence with the new merged evidence.

Once complete, we can eliminate any alleles that do not match the phased evidence.

##### 4. Recover common alleles

As a fail safe for phasing, any ‘common alleles’ with more than 0.1% population frequency are recovered. The frequencies of alleles are specified in a resource file and are derived from the Hartwig cohort.

Additionally, C*04:09N (the most common HLA allele with a frameshift variant) specifically is also rescued if the out of frame indel 6:31237115 CN>C is present.

##### 5. Detect HLA-Y presence

HLA-Y is a pseudogene that is highly similar to HLA-A and is not present in the human ref genome but is found in approximately 17% of the Hartwig cohort. The presence of HLA-Y can cause confusion in typing particularly in determining the HLA-A types. To detect HLA-Y, LILAC counts the number of fragments that can be assigned uniquely to one of the 3 known HLA-Y alleles and no other candidate alleles. If at least 1% of fragments align uniquely to HLA-Y then HLA-Y is considered to be present in the sample. If HLA-Y is found to be present ANY fragment which matches exactly to a HLA-Y allele (uniquely or shared with other alleles) are excluded from further analysis to prevent confusion with highly similar HLA-A alleles.

**6. Test for 2 digit types with unique evidence**

To further reduce the number of candidate alleles, If any 2-digit types are sufficiently unique (i.e. uniquely supported by at least 2% of fragments, they are required to contain at least one 4 digit type belonging to that 2 digit type in the evidence phase. If two 2 digit types from the same gene are found to be sufficiently unique all other alleles are discarded at this point. If more than two groups are found to be unique the 2 with the highest evidence are supported Any recovered alleles are also discarded at this point unless the 2 digit group has at least one fragment of unique support.

**7. Remove incomplete alleles with insufficient unique evidence**

Many alleles in the IMGT database are incomplete (ie contain ‘*’ characters), all of which are rare in population frequency. To prevent spurious matches to these wildcard containing alleles in the evidence phase, we eliminate unlikely candidates. Wildcard containing alleles are eliminated unless they contain at least 2 fragments support for the non wildcard sequence which do not support any remaining candidate allele with a complete sequence defined.

#### Evidence phase

In the evidence phase, LILAC evaluates all possible ‘complexes’ (ie. combinations) of remaining alleles that satisfy the following conditions

- There must be either 1 (homozygous) or 2 (heterozygous) alleles belonging to each gene
- At least 1 allele must match each uniquely supported 2 digit type

For each complex, LILAC counts the number of fragments that can be aligned exactly to at least one allele in the complex at all heterozygous locations. If a fragment has an amino acid which does not match ANY of the amino acid candidates at a heterozygous location, but has at least 1 nucleotide with base qual<min(medianBaseQuality,30), then the amino acid is deemed to match all amino acid candidates For exon boundaries only exact nucleotide matches are permitted. Any fragments that can be aligned to 2 or more alleles are apportioned equally between the alleles and counted as shared fragments.

Since many allele definitions have undetermined (‘wildcard’) sequences, particularly in exon 1 and exons 4-8, these require special treatment so that these wildcard alleles are neither unfairly favored or discriminated against in the fitting. To achieve this balance, fragments which don’t match an exact sequence in any candidate allele are dropped altogether from consideration such that random sequencing errors or other artifacts overlapping wildcard regions cannot contribute to the complex count for wildcard containing alleles, but any remaining fragments are considered to match an allele if they match all non wildcard sequences (ie. any amino acid is deemed a match to a wildcard).

Complexes are scored based on the total fragments that can be aligned to at least one allele in the complex, with a small penalty applied base on allele frequency in the population, a penalty for each allele included that was eliminated but subsequently recovered, and a bonus to boost scores of complexes with homozygous allele, and a penalty for solutions with wildcard characters which may cause spurious matches. The final score is given by:

```
Complex Score = AlignedFragments + FreqPenalty + HomBonus + RecoveryPenalty +
Wildcard penalty
where
   FreqPenalty = 0.0015 * SUM[max(log10(Frequency),1e-4) * AlignedFragments]
   HomBonus = 0.0045 * (# of Homozygous alleles) * Fragments
   RecoveryPenalty = 0.005 * (# of Recovered alleles) * Fragments
   WildcardPenalty = 0.000015 * (# of wildcard characters in alleles) * Fragments
```

If 2 complexes are precisely equally scored, then the solution with the lowest alphabetical 4 digit allele types is chosen.

The matching unique, apportioned shared, and wildcard (in rare cases where the full allele is not present in the IMGT/HLA database) support for each allele is recorded in both tumor and normal.

As a further performance optimisation, if there are predicted to be more than 1 million complexes, then the evidence phase is first performed individually for complexes of 2 alleles per gene to find the top candidates for each of HLA-A, HLA-B & HLA-C. LILAC retains only the top 5 pairs including each individual allele candidate and then chooses the first 10 unique alleles appearing in the ranked list of pairs, with any common alleles also retained. The evidence phase is then subsequently run using this reduced set of candidate alleles

#### Tumor and RNA status of alleles

##### Tumor allele specific copy number

LILAC optionally accepts a tumor .bam file and a gene copy number file (produced by PURPLE) which contains the minimum copy number and minimum minor allele copy number of each gene. If a tumor sample is provided, then fragments are counted for each allele in the determined type. For each of HLA-A, HLA-B & HLA-C, the minor allele copy number is assigned to the allele with the lowest ratio of supporting fragments for the allele in the tumor compared to the normal. The other allele for each is assigned the implied major allele copy number from the gene copy number file. If a gene is homozygous present in the normal sample, then the minor and major allele copy numbers are arbitrarily assigned in the tumor.

##### Somatic variant assignment to alleles

LILAC optionally accepts a VCF input of somatic small indel and point mutations (called by Sage in the Hartwig pipeline) and can assign somatic variants to the specific allele which is damaged.

LILAC gathers the set of variants from the vcf (filter = “PASS”) that overlap either a coding or canonical splice region in any of HLA-A, HLA-B and HLA-C and finds all fragments that contain that variant. LILAC assigns aportions the fragment to the allele which matches the fragment at all heterozygous locations after excluding any somatic variants from the fragment. The allele with the highest matching fragment count is determined to contain the somatic variant. If the variant is assigned to 2 alleles with identical weight it is assigned with 0.5 weight to each. In the case of homozygous alleles, the variant is assigned arbitrarily to the 1st allele.

##### RNA ‘expression’ of alleles

LILAC also optionally accepts a RNA bam. As per the fragments in the bam are counted for each allele in the determined type. This can be interpreted as a proxy for allele specific expression.

#### QC metrics and PON

LILAC produces a comprehensive set of QC metrics and provides warning statuses if the typing confidence may be diminished. The full list of possible QC warnings is

| Warning | Description |
|---|---|
| WARN_UNMATCHED_TYPE | all A, B or C types eliminated |
| WARN_UNMATCHED_INDEL | A novel (non PON) indel is present that was not fit to any allele supported by at least 0.5% of fitted fragments. |

| | |
|---|---|
| WARN_UNMATCHED_HAPLOTYPE | A novel (non PON) haplotype is present that was not fit to any allele supported by at least 1% of fitted fragments. |
| WARN_UNMATCHED_AMINO_ACID | A novel (non PON) amino acid is present that was not fit to any allele supported by at least 1% of fitted fragments. |
| WARN_UNMATCHED_SOMATIC_VARIANT | A somatic variant from the input vcf could not be assigned to any allele |
| WARN_LOW_BASE_QUALITY | Median base quality < 25 |
| WARN_LOW_COVERAGE | More than 50 distinct bases of HLA-A, HLA-B and HLA-C combined have less than 10 coverage. |
| FAIL_LOW_COVERAGE | More than 200 distinct bases of HLA-A, HLA-B and HLA-C combined have less than 10 coverage. |

In general the warnings may indicate one of either a novel germline variant/allele, presence of an unusual pseudogene type or an incorrectly typed sample.

In the case of unmatched haplotypes and indels, we find that there are a number of recurrent artifacts found across many samples in our cohort. We therefore 'PON' filter the following indels and haplotypes prior to calculating QC metrics for the fit

**INDEL PON**

| **37_Position** | **38_Position** | **Variant** | **Curation** |
|---|---|---|---|
| 29912028 | 29944251 | AN>A | Recurrent HLA-H artefact |

| | | | |
|---|---|---|---|
| 29911899 | 29944122 | A>ACC | Recurrent HLA-Y artefact |
| 29911318 | 29943541 | CN>C | Recurrent artefact |
| 31324524 | 31356747 | GNN><br>G | Phased alignment issue |
| 31324528 | 31356751 | G>GGT | Phased alignment issue |
| 29910716 | 29942939 | CN>C | Recurrent artefact |
| 29910657 | 29942880 | G>GT | Recurrent artefact |
| 29912392 | 29944615 | A>AGG | Recurrent artefact |
| 29911899 | 29944122 | A>AC | Recurrent artefact |

**HAPLOTYPE PON**

| Start | Haplotype | Curation |
|---|---|---|
| 1 | AVVAPRTLLLLLSGALALTQTWAG | HLA-Y |
| 25 | SHSMRYFSTSVSRPGSGEPRFIAVGYVDDTQF<br>VRFDSDAASQRMEPRAPWMEQEEPEYWDRQ<br>TEISKTNAQIDLESLRIALRYYNQSED | HLA-Y |
| 47 | AVGYVDDTQFVRFDSDAASQRMEPRAPWMEQ<br>EEPEYWDRQTQISKTNAQIDLESLRIALR | HLA-Y |
| 96 | IDLESLRIALRYYNQSEA | HLA-Y |
| 117 | TIQRMSGCDVGSDGRFLRGYRQDAYDGKDYIA<br>LNEDLRSWTAADMAAQITQRKWEAARQAEQL<br>RAYLEGECMEWLRRYLENGKETLQRT | HLA-Y |

| | | |
|---|---|---|
| 206 | DAPKTHMTHHAVSDNEATLRCXALSFYPAEITLTWQR | HLA-Y |
| 206 | DPPKTHMTHYPISDHEATLRCWALG | HLA-H |
| 206 | DPPKTHMTHHPISDHEATLRCWALGFYPAEITLTWQRDGEDQTQDTELVETRPAGDG | HLA-H |
| 206 | DPPKTHMTHHPISDHEATLRCWALSFYPAEITLTWQRDGEDQTQDTELVETRPAGDG | Likely HLA-Y |
| 242 | RDGEDQTQDTELVETRPAGDGTFQKWASVVVPSGQEQRYTCHVQHEGLPKPLTLRWE | Likely HLA-Y |
| 262 | GIFQKWAAVVVPSGEEQRYTCHVQHEGLPKPLTLRWE | HLA-Y |
| 298 | EPSSQPTIPIVGILAGLVLFGAVIAGAVVAAVMWRRKS | Likely HLA-Y |
| 299 | PSSHPTIPIVGILAGLVLFGAVIAGAVVAAVMWRRKS | HLA-Y |
| 299 | IPNLGIVSGPAVLAVLAVLAVLAV | Alignment issue with C*17 indel |
| 337 | DRKGGSYSQAAS | HLA-Y or HLA-H |
| 351 | IAQGSDVSLTAC | HLA-Y |

#### Output

The following files are written:

#### Solution Summary

| Field | Description |
|---|---|
| Allele | Allele ID |
| RefTotal | Total assigned fragments from reference BAM |
| RefUnique | Fragments uniquely assigned to allele |
| RefShared | Fragments assigned to allele and others in this solution |
| RefWild | Fragments matched to a wildcard allele |
| TumorCopyNumber | Copy number from Tumor/Ref fragment ratio and Purple copy number |
| TumorTotal | As above for tumor BAM |
| TumorUnique | As above for tumor BAM |
| TumorShared | As above for tumor BAM |
| TumorWild | As above for tumor BAM |
| RnaTotal | As above for RNA BAM |
| RnaUnique | As above for RNA BAM |
| RnaShared | As above for RNA BAM |
| RnaWild | As above for RNA BAM |
| SomaticMissense | Matched missense variants |

| | |
|---|---|
| SomaticNonsenseOrFrameshift | Matched nonsense or frameshift variants |
| SomaticSplice | Matched splice variants |
| SomaticSynonymous | Matched synonymous variants |
| SomaticInframeIndel | Matched inframe indels |

#### QC Metrics

| Field | Description |
|---|---|
| Status | Either PASS or 1 or more warnings (see below for warning descriptions) |
| HlaY | HLA-Y allele detected if present, else ‘NONE’ |
| ScoreMargin | Difference in score to second-top solution |
| NextSolutionAlleles | Allele difference in second-top solution |
| MedianBaseQuality | Median base quality across all coding bases from all fragments |
| DiscardedIndels | Discarded fragments due to unknown INDELs |
| DiscardedIndelMaxFrags | Maximum fragment support for an indel detected but not present in any known allele. |
| DiscardedAlignmentFragments | Fragments discarded because 1 read aligns more than 1000 bases from a HLA A,B or C gene |

| | |
|---|---|
| A_LowCoverageBases | Number of bases with less than 15-depth coverage across all coding bases, also for B and C genes |
| ATypes | # of distinct HLA-A alleles fitted (0,1 or 2) |
| BTypes | # of distinct HLA-B alleles fitted (0,1 or 2) |
| CTypes | # of distinct HLA-C alleles fitted (0,1 or 2) |
| TotalFragments | Total of fragments overlapping a coding base of HLA-A, HLA-B or HLA-C with MAPQ >=1 |
| FittedFragments | Total fragments assigned to fitted alleles (ie. exact or wildcard match at every heterozygous location) |
| UnmatchedFragments | Fragment that do not match any allele exactly in solution at all heterozygous location (eg. due to sequencing error or mapping error) |
| UninformativeFragments | Fragments that do not overlap any heterozygous location considered in evidence phase |
| HlaYFragments | Fragments excluded from fit due to allocation to assignment to HLA-Y. |
| PercentUnique | Percentage of fitted fragments that are uniquely assigned to 1 allele |
| PercentShared | Percentage of fitted fragments allocated across multiple alleles |
| PercentWildcard | Percentage of fitted fragments uniquely assigned to wildcard regions |

| | |
|---|---|
| UnusedAminoAcids | # of amino acids detected with at least 3 fragments support but not present in final allele solution set (excluding PON filtered and regions overlapping wildcards in selected alleles) |
| UnusedAminoAcidMaxFrags | Maximum fragments support for an unmatched amino acid not present in the final allele solution set |
| UnusedHaplotypes | # of haplotypes phased with at least 3 fragments support but not present in final allele solution set (excluding PON filtered and regions overlapping wildcards in selected alleles) |
| UnusedHaplotypeMaxFrags | Maximum support for an unmatched haplotype not present in final allele solution set |
| SomaticVariantsMatched | Somatic variants supported by solution allele |
| SomaticVariantsUnmatched | Somatic variants not supported by solution allele |

**Additional output files**

| File | Description |
|---|---|
| SAMPLE_ID.lilac.somatic.vcf.gz | Annotation of supplied somatic variants if somatic VCF provided with assigned allele |
| SAMPLE_ID.fragments.csv | Read details for all BAM fragments |
| SAMPLE_ID.candidate.coverage.csv | Coverage for all candidate solutions within X% of the top solution's score |

| | |
|---|---|
| SAMPLE_ID.candidate.fragments.csv | Allocation of each fragment to one or more solutions and which alleles they support |
| SAMPLE_ID.HLA-A.aminoacids.txt | Fragment support for each amino acid by HLA gene |
| SAMPLE_ID.HLA-A.nucleotides.txt | Fragment support for each nucleotide by HLA gene |
| SAMPLE_ID.candidates.aminoacids.txt | Fragment support for amino acids in the candidate alleles |
| SAMPLE_ID.candidates.nucleotides.txt | Fragment support for nucleotides in the candidate alleles |

The log file contains additional detailed information about the fit including details of all unmatched haplotypes and indels.
