## Supplementary Note 2 for "Genetic immune escape landscape in primary and metastatic cancer"

### Neo pipeline

#### Overview

The neoepitope pipeline (Neo) works in 2 main steps to form a comprehensive set of neoepitope and neopeptide predictions from our DNA pipeline output:

- Determination of all novel peptides (i.e., neopeptides) from all point mutations, small indels and gene fusions.
- Calculation of allele specific binding affinity and presentation scores using a novel binding affinity prediction algorithm.

Although we annotate with expression information from RNA (where available), the neoepitope predictions are currently based solely on mutations found in DNA. Hence we specifically ignore RNA events such as circular RNA, RNA editing, endogenous retroviruses and alternative splicing as we are unable to determine if these are tumor specific and hence will make neoepitopes. High confidence fusions detected in RNA but not found in DNA are also currently ignored (see future improvements). We also acknowledge that we miss protein level events including non-canonical reading frames, post translational amino acid modifications & proteasomal peptide splicing.

##### Inputs

Neo use the following inputs from the Hartwig pipeline

- **Somatic variants:** PURPLE somatic .vcf
- **Structural variants:** LINX candidate fusion neoepitopes (a new file produced by LINX to output all candidate neoepitope files).
- **HLA typing:** LILAC output (or from alternative methods as long as they are formatted appropriately).

Where RNA is available additional annotations are added, an effective TPM of each neoepitope is also estimated based on the following inputs

- **Gene Expression:** Isofox (https://github.com/hartwigmedical/hmftools/tree/master/isofox) transcript expression
- **Neo-epitope fragment support:** RNA .bam

For presentation and immunogenicity predictions, Neo also uses a number of resource files, that are pre-calculated from external resources (including IEDB[1], the HLAthena[2] publication and the IPD-IMGT/HLA[3] database), specifically:

| File Name | Purpose | Allele specific generation |
|---|---|---|
| isofox.transcript_medians.csv | Median TPM per gene per cancer type and pan-cancer | No |
| IEDB_known_immungoenic_peptides.csv | IEDB known immunogenic peptides | No |
| neo_TPM_likelihood_disribution.csv | Mean pan-allele presentation likelihood by decile and TPM bucket, precalculated from HLAthena data | No |
| neo_train_flank_pos_weight.csv | Pre-calculated pan-allele flank position counts and weight matrix | no |
| neo_train_pos_weight.csv | Pre-calculated allele specific position counts and weight matrix | Yes |
| neo_train_length_specific_score_rand_dist.csv | Pre-calculated percentile ranks of length presentation likelihood for random peptides | Yes |
| neo_train_length_specific_rank_to_likelihood.csv | Pre-calculated conversion from length specific likelihood rank to likelihood | Yes |
| neo_train_likelihood_rand_dist.csv | Pre-calculated percentile ranks of presentation likelihood for random peptides | Yes |
| neo_train_exp_likelihood_rand_dist.csv | Pre-calculated percentile ranks of expression presentation likelihood for random peptides | Yes |

### Workflow overview

There are 2 key steps in the Neo pipeline:

1. Prediction of candidate neoepitopes from somatic variants
2. Estimation of presentation likelihood of each candidate pHLA ({peptide,allele})

##### 1. Candidate neoepitopes

We search for potential neoepitopes for point mutations and structural variants which meet the following criteria

| Type | Criteria |
|---|---|
| Point mutations | • Filter = 'PASS'<br>• Coding effect in (missense, frameshift, in-frame indel and stop lost) |
| Fusions (intergenic or intragenic) | Rules as per LINX fusion calls with the following exceptions:<br>• The 5' partner breakend for neo-epitopes MUST be in the coding region (exonic or intronic)<br>• No restriction applies on the coding context for the 3' breakend for neoepitopes<br>• Fusions that are predicted to be terminated in the 5' or 3'are not considered for neo-epitopes<br>• The 3' transcript biotype for neo-epitope must not be 'nonsense mediated decay' |

Subject to the criteria above, all transcripts (or combination of transcripts in the case of fusions) are considered as candidate neoepitopes. Where 2 transcripts (or transcript combinations) lead to either the same amino acid sequence or the amino acid sequence of one transcript forms a subset of another, the transcripts are merged to form a single neoepitope. For each unique neoepitope, Neo outputs the amino acid (AA) sequence string broken up into 'upstream', 'novel' and 'downstream' segments as follows:

| Field | Description |
|---|---|
| NeId | Unique Id for neoepitope |
| Variant type | One of: {MISSENSE, INFRAME, OUT_OF_FRAME_FUSION, INFRAME_FUSION, FRAMESHIFT} |
| VariantInfo | Unique identifier for variant<br>For point mutations = <chr>:<Position>:<ref>:<alt><br>For SV = <chrUp>:<posUp>:<orientUp>-<chrDown>:<posDown>:<orientDown> |
| GeneNameUp | Gene name for the upstream part of the neoepitope |
| GeneNameDown | Gene name for the downstream part of the neoepitope |
| UpstreamAA | Section of the neoepitope that matches the upstream transcript |
| DownstreamAA | Section of the neoepitope that matches the downstream transcript (if any) |
| NovelAA | Novel section of the neoepitope (if any) |
| WildtypeAA | Wildtype AA sequence for agretopicity calculation (missense variants only) |
| UpTranscripts | List of transcripts in the up gene that support the neoepitope |
| DownTranscripts | All unique transcripts on the up gene that support the neoepitope |

Note that the precise definition of the novel segment and upstream and downstream AA depend on the type of event. The exact rules are outlined in the table below:

| Variant Type | Novel Segment | Upstream flank | Downstream flank |
|---|---|---|---|
| Missense SNV/MNV* | Ref->Alt AA(s) | Up to 16 AA limited by start codon | Up to 16 AA limited by stop codon |
| Inframe* | If conservative inframe, inserted AA only else also use flanking disrupted AA on each end | Up to 16 AA limited by start codon | Up to 16 AA limited by stop codon |
| Stop_lost / frameshift* | All downstream AA until new stop codon reached | Up to 15 AA limited by start codon | NA |
| Inframe fusion (Phase=0) | NA*** | Up to 16 AA limited by start codon | Up to 16 AA limited by stop codon |
| Inframe fusion (Phase = {1,2}) | Mixed transcript AA*** | Up to 16 AA limited by start codon | Up to 16 AA limited by stop codon |
| Out of frame coding to coding or coding to non-coding fusion | Possible mixed transcript AA + all downstream AA until new stop codon | Up to 16 AA limited by start codon | NA** |

| | |
|---|---|
| | reached** |

* Where multiple somatic variants are phased within 17 AA, include entire intermediate section as novel AA
** For coding to 5’UTR fusions, if a start codon is reached prior to a novel stop codon and is ‘inframe’, the novel segment should be limited to the region up to the new stop codon with the downstream flank set as the first 17 AA of the 3’ partner.
*** For exonic-exonic fusions, include any inserted sequence

Neo further annotates each of the candidate neoepitopes with TPM and direct RNA fragment support for the novel amino acid sequence. TPM per transcript is sourced from Isofox (https://github.com/hartwigmedical/hmftools/tree/master/isofox) if RNASeq is available or if not available is estimated as the median of the cancer type (or full cohort where cancer type is not known). Neo also reanalyses the RNA .bam to count the RNA depth at the location of the variant that caused the neoepitope and the direct RNA fragment support for the neoepitope (defined as matching precisely the 1st novel AA and 5 bases either side).

#### 2. pHLA presentation likelihood

Using the identified neoepitopes, we determine all candidate pHLA ({peptide,allele}) combinations that may be presented by the cell. For each candidate neoepitope, we consider all peptides between 8 and 12 length which either overlap the novel amino acid sequence or overlap both the upstream and downstream amino acid sequence.

For each pHLA we determine a presentation likelihood (see below Novel pHLA presentation likelihood),and a number of immunogenic features:

| Field | Description |
|---|---|
| NeId | Identifier of neoepitope from which peptide was obtained |
| VariantType | See neoepitope |
| VariantInfo | See neoepitope |
| GeneNameUp | See neoepitope |
| GeneNameDown | See neoepitope |
| HlaAllele | MHC class 1 allele for which presentation likelihood have been calculated |
| HlaAlleleDisruption | "LOH','SOMATIC' or 'NONE' |
| Peptide | Amino acid sequence of peptide |
| n_flank | 5’ flanking amino acid sequence (up to 3 AA) |
| c_flank | 3’ flanking amino acid sequence (up to 3 AA) |
| LengthScore | Raw score for {peptide,allele} pairing |
| LengthScoreRank | Presentation percentile by length and allele |
| PresentationLikelihood | Relative likelihood of presentation by allele |

| | |
|---|---|
| PresentationRank | Presentation percentile by allele |
| ExpAdjLikelihood | Expression adjusted likelihood of presentation by allele |
| ExpAdjRank | Expression adjusted presentation percentile by allele |
| WildtypePeptide | Wildtype peptide (for missense only, else NA) |
| NearestPeptideProteome | Peptide with lowest BLOSSUM62 distance in human proteome |
| NearestPeptideBinder | Binding Peptide (presentationRank <0.01 with lowest BLOSSUM62 distance in human proteome |
| DisToWildType | BLOSSUM62 distance to wildtype peptide (for missense only, else NA) |
| DisToProteome | Lowest BLOSSUM62 distance to any peptide of same length in human proteome |
| DisToBinder | Lowest BLOSSUM62 distance to any binding peptide of same length in human proteome with presentation rank <0.01 |
| DisToKnownImmunogenic | Lowest BLOSSUM62 distance to any peptide of same length to any known immunogenic peptide from IEDB |
| MutatedPosition | Anchor positions affect binding / allele preference relative to W/T whereas non anchor positions have similar binding but may impact immunogenicity. NA for frameshift |

### Novel pHLA presentation likelihood algorithm

#### Conventions regarding positions, peptide lengths and HLA allele binding motifs

##### Position Independence

Our model assumes that the peptides at each position each have an independent impact on binding and that no correlated effects apply (an assumption held almost universally across binding prediction tools). In fact, a preliminary internal test showed that this assumption holds true.

##### Peptide length mappings

Neo supports 8-12 length kmers. Our model assumes that anchor (2nd and last peptide position) positions and their surrounding positions have high similarity across peptide length with central peptides variable and relatively less important for binding. We therefore convert all peptides to a 12mer, with the following padding conventions for shorter peptides.

| | | | | | | | | | | | | |
|---|---|---|---|---|---|---|---|---|---|---|---|---|
| **12mer** | 0 | 1 | 2 | 3 | 4 | 5 | 6 | 7 | 8 | 9 | 10 | 11 |
| **11mer** | 0 | 1 | 2 | 3 | 4 | X | 5 | 6 | 7 | 8 | 9 | 10 |
| **10mer** | 0 | 1 | 2 | 3 | 4 | X | X | 5 | 6 | 7 | 8 | 9 |
| **9mer** | 0 | 1 | 2 | 3 | 4 | X | X | X | 5 | 6 | 7 | 8 |
| **8mer** | 0 | 1 | 2 | 3 | 4 | X | X | X | X | 5 | 6 | 7 |

##### Flanking sequences

The flanking amino acids upstream and downstream are known to impact cleavage and proteasomal processing. To capture these impacts we include 3 upstream amino acids (U3,U2,U1) and 3 downstream amino acids (D1,D2,D3) in the model. The enrichment/depletion of amino acids at these positions globally (including 'X' where the flanking sequences are beyond the start or end of an allele) is included in the peptide score.

##### Binding Motifs

We utilize the same assumptions as NetMHCpan[4] which is that only proximate (within 4 angstroms across a representative set of HLA-A/ HLA-B structures) polymorphic residues may affect binding, which yields 34 distinct positions with the following specificities.

| Position | Proximate Polymorphic Amino Acid Residues |
|---|---|
| 0 | 31,83,86,87,90,183,187,191,195 |

| | |
|---|---|
| 1 | 31,33,48,69,86,87,90,91,94,123,183 |
| 2 | 94,121,123,138,180,183 |
| 3 | 90,182,183,187 |
| 4 | 93,94,182 |
| 5 | 93,94,97,98,121,180 |
| 6 | 93,97,121,138,171,174,176,180 |
| 7 | 97,100,101,171 |
| 8 | 98,101,104,105,108,119,121,140,142,167,171 |

In general we observe that positions with identical binding motifs observe highly similar amino acid weight distributions in mass spectrometry observations.

##### Binding Motif similarity

We assume that binding motifs with similarity tend to have similar bindings. We estimate binding motif similarity by summing the log likelihoods from the BLOSUM62 substitution matrix across all binding positions.

#### Training dataset

We have curated IEDB and the literature for high quality unbiased monoallelic mass spectrometry results assessing that i) the datasets were monoallelic and ii) whether the study did not appear to contain an empirically high rate of likely false positive results. Binding affinity results are not used in the training data due to their inherent experimental selection bias. Overall we identified 20 studies (all included in IEDB) with 413,000 MS observations across 103 alleles to be used in our training set. For 8 specific alleles which had no mono-allelic results, but where there was sufficient high quality multi-allelic data (specifically A*69:01,B*35:08,B*41:01,A*26:08,C*15:05',B*44:09,B*44:27 and B*44:28) we included the results from 4 additional studies.

Finally, we also included monoallelic data from the recent Pyke et al.[5] (hereafter also referred to as Sherpa dataset) study which has results from 25 monoallelic cell lines including 6 additional cell lines not represented in the IEDB data. To eliminate potential experimental artifacts, the Sherpa dataset was also filtered to include only peptides that were found to be bound to 2 or less distinct alleles.

Any {peptide;allele} observations found in more than 1 dataset may be counted multiple times towards position matrices. We also have determined the 3 bases flanking both upstream and downstream for each peptide where possible by matching the peptides to the reference proteome. 10% of the full data is held back as a validation dataset (see Validation section).

#### Construction of Position Weight Matrix (PWM)

A position weight matrix is constructed per allele per peptide length. To deal with sparsity of data for specific alleles and peptide lengths, we use the following principles to learn from other peptide lengths and alleles:

1. Peptide lengths may learn from other lengths when they lack sufficient allele and length specific data. The more similar the peptide length, the higher the weighting

2. Alleles may learn from other alleles with identical or similar binding motifs for that position when they lack sufficient allele specific data. The more similar the binding motif, the higher the weighting

This is implemented in a 2 step process. First we consolidate all specific length counts into a single length weighted count (LWCount) for the tested peptide length for each allele using the following formula:

LWCount(A,L,P,AA) = Count(A,L,P,AA) + LWF * SUM(l<>L) [Count(A,l,P,AA) /abs(L-l)] * maxLW / max( LWF * SUM(l<>L)[Count(A,l,P) /abs(L-l)],maxLW)

Then, using the motif m of the tested allele, we consolidate all matching and similar binding motif observations from other alleles to obtain a total weighted count (WCount) using the following formula:

WCount(A,L,P,AA) = LWCount(A,L,P,AA) + MWF* SUM(a<>A) [ LWCount(a,L,P,AA) * (2^(LogSim(m,M)) / MAX(i=all motifs)[2^(LogSim(i,M))]] * maxMW / max(MWF * SUM(a<>A) [ LWCount(a,L,P) * (2^(LogSim(m,M)) / MAX(i=all motifs)[2^(LogSim(i,M))]],maxMW)

where:

- A=allele
- L= length
- P= Peptide position
- AA = amino acid
- M = binding motif of allele A at position P
- LWF = length weight factor (<=1; default = 0.25)
- MWF = motif weight factor (<=1; default = 0.25)
- maxLW = max length weight (default = 200)
- maxMW = motif weight factor (default = 200)
- Obs(a,l) = observations of peptides of binding allele a and peptide length l
- LogSim(a,b) = Blosum62 log similarity of motifs a and b summed over all motif positions

The output of this is a final weighted position weight matrix for each allele and length.

#### Scoring and ranking per allele per peptide length

Each peptide is scored based on the positional amino acid frequencies relative to the amino acid frequency in the proteome:

**PeptideScore = Sum[Log2(max(P(x,i),0.005)/Q(x))]**

where
- P(x,i) = % weight of amino acid = x at position = i (note for C we use 3*weight to correct for MS bias)
- Q(x) = frequency of amino acid = x in the proteome

An additional flank score based on a pan allele flanking PWM is calculated as follows:

**FlankScore = Sum(i=U3-U1,D1-D3)[Log2(P(x,i)/Q(x))]**

As has been noted previously by other groups, we do observe enrichment when U1 = 'M*' (ie the first amino acid of the coding sequence) or D1 = 'X' the stop codon of a transcript, but not for the other bases in the flanks. Hence for the U1 and D1 base we set the PWM score to be the observed enrichment for 'M*' (approximately 2 fold greater) and 'X' (approximately 4 fold greater) respectively. For the other 2 flanking bases on either side we observe no further enrichment in 'M*' or 'X' and hence set the PWM score simply to 0 (no enrichment or depletion) if they overlap the start or end of the transcript.

The total score is then set to:

**TotalScore = PeptideScore + FlankScore**

The score is converted to a binding rank percentile (per allele per peptide length) by comparing to the percentile scores compared to scores for 100,000 peptides of the same length randomly from the proteome (note that random peptides are excluded if they are found to be binders already in the MS results). Where flanks are available the rank is given relative to the total score distribution and where not available relative to just the peptide score distribution.

##### Relative presentation likelihood and ranking per allele (pan peptide length)

To assess the relative likelihood of presenting peptides of different lengths for a particular allele given the binding ranks, we determine the relative density of mass spectrometry observations in our training set per peptide length per binding rank percentile bucket. The relative likelihood is calculated as:

**RelPresentationLikelihood = MSobs(A,Rb,L)/Size(Rb)/SUM(Rb,L)[MSobs(A,Rb,L)/ Size(Rb)]**

Where
- A = Allele

- L = Peptide Length
- Rb = Rank Bucket. Exponential buckets in powers of 2 are used to reflect the relative importance of the very low binding ranks: {0.00005,0.0001,0.0002,0.0004,...,0.8192}

To deal with sparse MS data, particularly for non-9mers, the density of a given ranking bucket is set to be at least as high as any lower ranked bucket. Furthermore, so that we can always rank the higher buckets regardless of the number of observations, any bucket with 0 observations is padded with observations of min(totalObservationsPerAllele/1000,0.25) for bucketed rank < 1%, or half the observations of the preceding ranked bucket where the bucketed rank > 1%

The output of this algorithm is a set of weights per bucket per peptide length reflecting the relative likelihood of an observation from that bucket being presented on the surface in that cell. For individual peptides we can predict an exact relative likelihood by interpolating the rank between the bucketed values

An example of the output for A*29:02 is shown below indicating that a 9mer with 0.00005 rank is ~28x (ie 0.3007/0.0116) more likely to be presented than an 8mer with the same rank and ~6x (ie 0.3007/0.0520) more likely to be presented as a 9mer with a rank of 0.0004.

| | Bucketed Rank | MS Obs by allele by length by ranking | | | | | Relative Presentation likelihood | | | | |
|---|---|---|---|---|---|---|---|---|---|---|---|
| | | 8 | 9 | 10 | 11 | 12 | 8 | 9 | 10 | 11 | 12 |
| A2902 | 0.00005 | 14 | 364 | 100 | 89 | 36 | 0.0116 | 0.3007 | 0.0826 | 0.0735 | 0.0297 |
| | 0.0001 | 8 | 402 | 67 | 61 | 26 | 0.0033 | 0.1661 | 0.0277 | 0.0252 | 0.0107 |
| | 0.0002 | 1 | 438 | 52 | 44 | 6 | 0.0009 | 0.0905 | 0.0125 | 0.0091 | 0.0014 |
| | 0.0004 | 9 | 504 | 121 | 50 | 14 | 0.0009 | 0.0520 | 0.0125 | 0.0052 | 0.0014 |
| | 0.0008 | 10 | 656 | 118 | 35 | 18 | 0.0006 | 0.0339 | 0.0061 | 0.0018 | 0.0009 |
| | 0.0016 | 23 | 691 | 146 | 41 | 19 | 0.0006 | 0.0178 | 0.0038 | 0.0011 | 0.0005 |
| | 0.0032 | 28 | 492 | 166 | 44 | 19 | 0.0004 | 0.0064 | 0.0021 | 0.0006 | 0.0002 |
| | 0.0064 | 28 | 382 | 99 | 38 | 7 | 0.0002 | 0.0025 | 0.0006 | 0.0002 | 0.0000 |
| | 0.0128 | 17 | 323 | 95 | 22 | 12 | 0.0001 | 0.0010 | 0.0003 | 0.0001 | 0.0000 |
| | 0.0256 | 20 | 166 | 55 | 11 | 1 | 0.0000 | 0.0003 | 0.0001 | 0.0000 | 0.0000 |
| | 0.0512 | 12 | 93 | 19 | 5 | 3 | 0.0000 | 0.0001 | 0.0000 | 0.0000 | 0.0000 |

The relative presentation likelihood is calculated for each 4 digit and 2 digit allele with more than 200 MS observations in the training data as well globally for HLA-A, HLA-B and HLA-C. Any 4 digit allele which does not have sufficient MS observations is assigned the likelihoods of the 2 digit allele if available or if not the likelihoods for the HLA gene as a whole.

A presentation percentile rank is calculated for each allele across all lengths by comparing the relative likelihood of the peptide compared to that of all negative decoy peptides (equally weighted across 8,9,10 and 11-mers to give best compatibility to other tool rankings).

### Expression adjusted presentation likelihood algorithm

#### Training data

For training the impact of expression on presentation likelihood we use the subset of the mass spectrometry training dataset included with the HLAthena publication[2] together with the matched TPM estimates provided with this publication for the B.721.221 cell line. For the purpose of the training the TPM of the gene is assumed to be the TPM of the transcript containing the epitope.

#### TPM adjusted likelihood rank

To determine the impact TPM expression has on presentation we compared the TPM of the MS identified peptides of strong predicted binders (LRank<0.1%) found to be presented in the HLAthena training data to all predicted strong binders from the proteome for the same alleles. For each $\log_2$ TPM bucket we calculate the proportion of pHLA combinations that are found to be presented. We find this to be a very strong relationship, ranging from a <<1% chance of presentation where TPM < 1 up to higher than 20% chance where TPM > 1000:

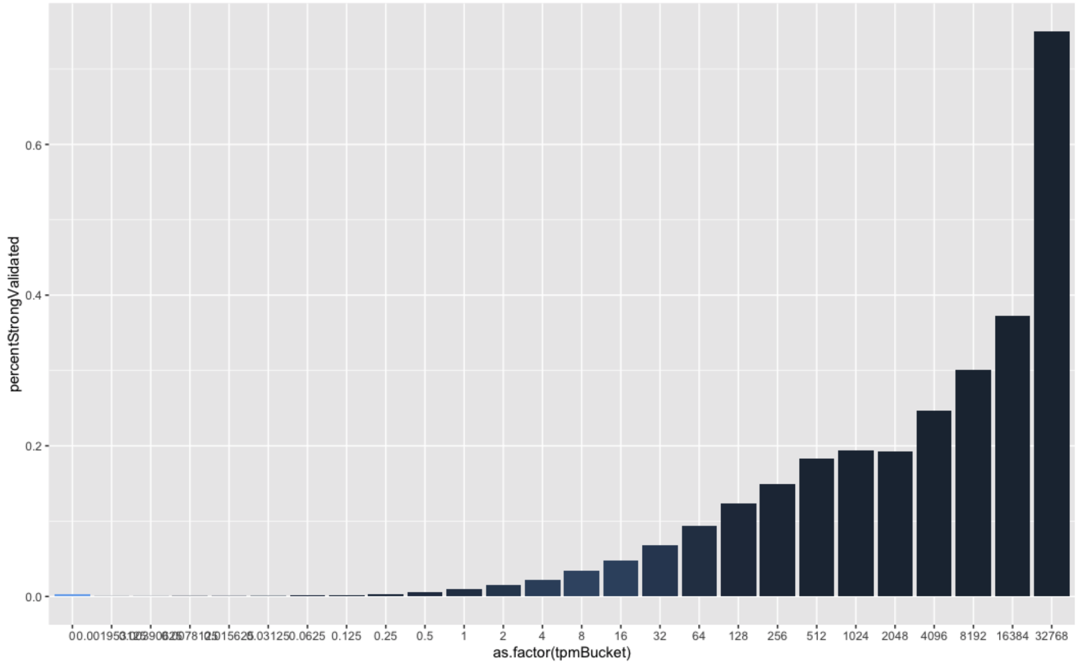

Using this observed rate of TPM we can calculate:

**ExpAdjLikelihood = PresentationLikelihood * TPMLikelihood / [PresentationLikelihood * TPMLikelihood + (1-PresentationLikelihood) * (1-TPMLikelihood)]**

We then calculate a new overall rank globally across all pHLA using the expression adjusted likelihood compared to the same randomly selected peptides.

### Neo pipeline validation

We assessed the performance of our neoepitope prioritization pipeline (Neo) by conducting four orthogonal validations:

1. Neo performance compared to MHCFlurry2.0 in a curated dataset of experimentally identified peptides derived from HLA monoallelic cell lines.
2. Neo ranking of random peptides.
3. Neo robustness by performing an allele leave-one-out experiment.
4. Neo expression adjusted model performance compared to the non-expression model.

For these validations, we assumed that the 100,000 random peptides used for the ranking likelihood calculation (after filtering any {peptide,allele} combinations included in the training data) were not binders. This is a conservative assumption as some of the highest ranked random peptides would certainly be expected to bind. Therefore, since only the top ~0.1% of peptides are expected to strongly bind, we mostly used the True Positive Rate (TPR) as the performance metric. This metric measures how well the model is able to rank known presented peptides (from a curated set) compared to the random set of 100,000 peptides. Other measurements, such as AUC measurements may be dominated by the relative performance of very weak predictions and are therefore not as informative in our endeavor.

#### 1. Neo performance compared to MHCFlurry2.0

The aim of this validation is to compare our pipeline ability to prioritize bona-fide peptides presented by the HLA-I in comparison to an outstanding open access tool such as MHCFlurry2.0[6]. To do so, we held-out 10% of the training set composed by the non-redundant union of experimentally identified peptides reported in IEDB and in other studies (see above Training set section). Hence, this 10% was not used in our training and will be used as a validation dataset to evaluate the True Positive Rate (TPR) across multiple rank thresholds compared to MHCFlurry2.0[6]. The TPR was estimated as the number of predicted peptides below the given threshold compared to the total number of peptides in the validation dataset. Moreover, to ensure comparability we use the presentation percentile of MHCFlurry2.0 and then recalculated a rank using 100k random peptides in the same manner we did for the Neo Likelihood ranks. This number deviates from the MHCFlurry Presentation rank which may have different assumptions about negatives

The average TPR was higher in Neo predictions compared to the predictions from MHCFlurry2.0 by relying in six ranking thresholds ($1e^{-06}$, $1e^{-05}$, $1e^{-04}$, $5e^{-04}$, $1e^{-03}$, $1e^{-02}$, $2e^{-02}$ and $1e^{-01}$, see Figure1 below). This trend was maininted when we split by peptide length from 8-kmers to 11-kmers (Figure 1). Nothewortly, the greatest differences in TPR were observed for 8-mers (average TPR Neo 0.48 compared to 0.32 of MHCFlurry2.0), while the difference for other peptide lengths was considerably lower. Taken together our results show that Neo sensitivity to rank true binders is slightly better than MHCFlurry2.0 at different percentile rank thresholds

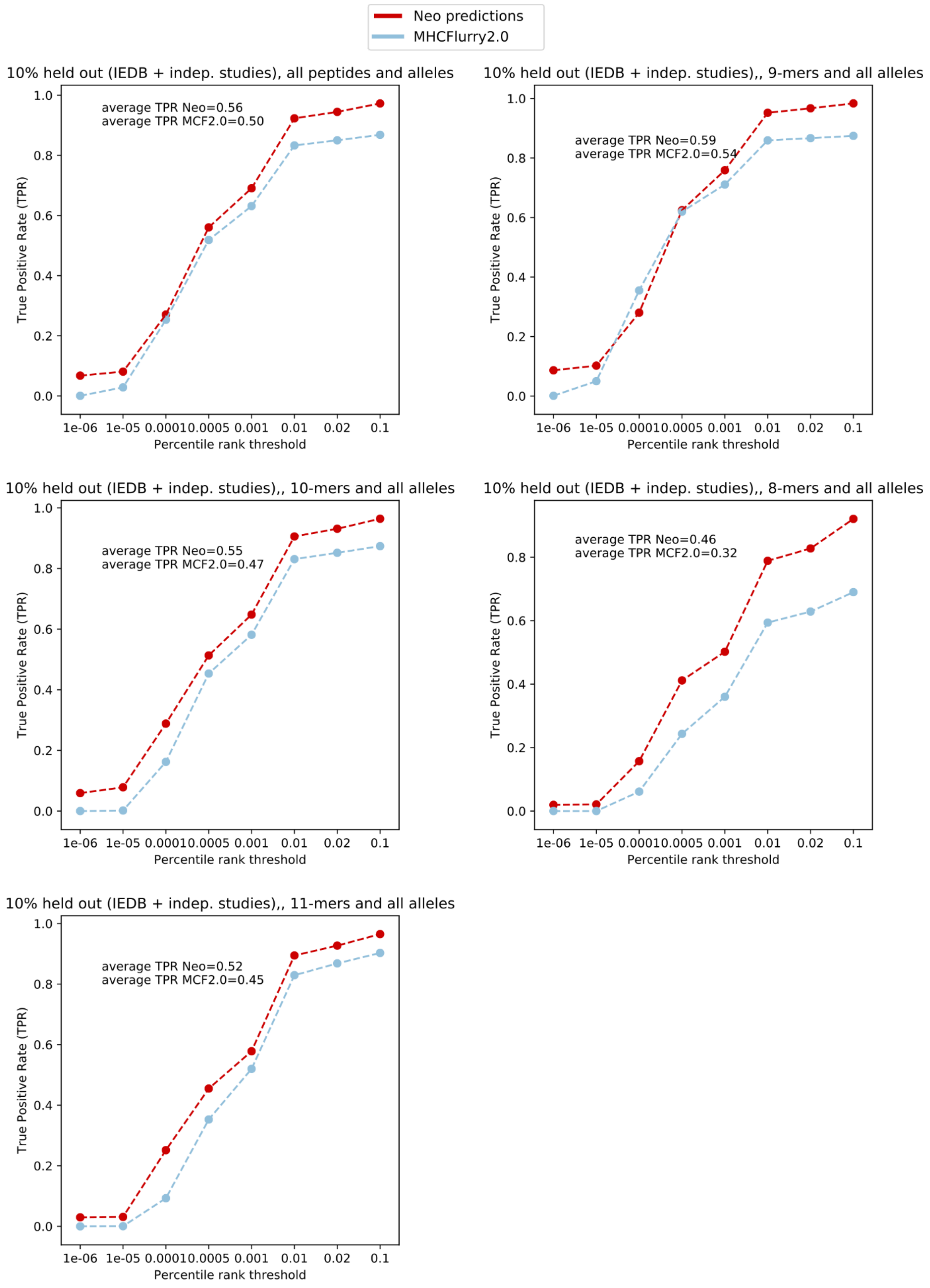

**Figure 1. Neo performance in a 10% held-out validation dataset compared to MHCFlurry2.0.** The x-axis represents the threshold of the ranking used as predicted

peptides. The y-axis represents the TPR at each threshold. Red and blue dots and lines represent Neo and MHCFlurry2.0 performance, respectively.

#### 2. Neo ranking of random peptides.

In the previous analysis we have shown Neo's sensitivity to rank true binders among the lowest percentile ranks. However, whether this is a unique feature of true binders or a systemic bias towards low percentile rankings is yet unclear. To address this, we evaluated Neo median percentile ranking in a set of 50k random peptides at five peptide lengths (from 8-12mers, 10k peptides per kmer length).

Reassuringly, we observed that the median allele percentile ranking distribution was very close to 0.5 (ie., percentile rank 50th) across all evaluated lengths and alleles (Figure 2). This results shows that observed percentile rankings for true binders are not the effect of a systemic bias towards low rankings and are thus the result of Neo's ability to discern between true binders are random peptides.

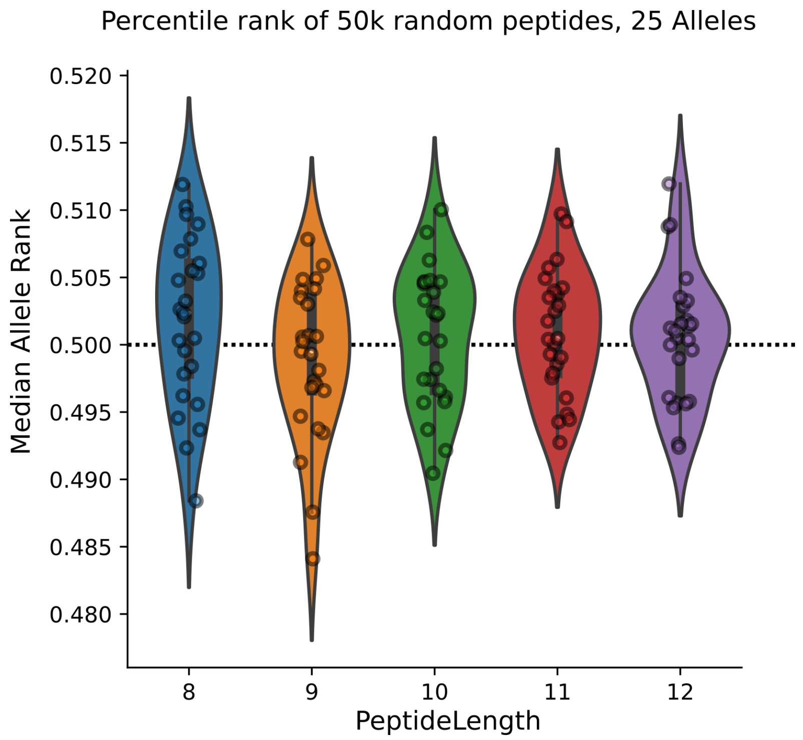

**Figure 2. Neo median percentile ranking in a set of 50,000 random peptides across 25 HLA-I alleles.** Each dot represents the median percentile rank across the evaluated random peptides for a particular allele and peptide length. Dashed horizontal line represents the 50th percentile ranking.

#### 3. Neo robustness by performing an allele leave-one-out experiment.

We aimed to assess how well our model is able to rank the presentation of peptides for an HLA allele that lacks representation in the training set. To do so we performed the following steps:

I. For each of the 116 HLA alleles with sufficient representation in the training dataset we iteratively removed the peptides from the curated dataset and re-trained the full model (i.e., leave-one-out experiment).
II. We computed the TPR at a 0.02 ranking threshold on the curated peptides for that allele (i.e., number of ranked peptides with a ranking likelihood below that threshold compared to the total number of peptides for that allele in the original training set).
III. We compared the leave-one-out TPR to the original TPR by training with the full dataset. The percentage of TPR decrease (%TPR loss) measures the ability of our model to generalize predictions for HLA alleles that lack representation. Alleles with high %TPR loss are those for which the model can not reliably make predictions without the training data. Conversely, HLA alleles with low %TPR loss are those for which the model can find accurate predictions by generalizing the predictions from other alleles with available training data.

We observed an average %TPR loss of 4.4% ± 6% std. (Figure 3). Only 15 HLA alleles (~12% of the 116 screened alleles) show a %TPR loss greater than 10% (Figure 3), highlighting the capacity of Neo pipeline to generalize predictions for alleles lacking representation in the training set. Certain alleles such as B*08:01, B*15:03 or A*30:01 displayed a very high %TPR loss suggesting that they may have a unique binding preference that can not be interpreted from neighbor HLA alleles (Figure 4). Another plausible explanation is that the 34 HLA amino acids selected for our model are unable to capture the binding preference of these alleles and additional amino acids are thus needed.

When splitting by peptide length, we observed that the most consistent predictions were for 9-mers, likely due to the higher representativeness of these peptides in the training dataset. Other k-mers had greater average %TPR loss.

Taken together our results show that, in general, our pipeline is able to accurately rank peptides for HLA alleles lacking representation in the training set and that our learn-from-others approach is a robust strategy to prioritize peptides presented by the HLA complex.

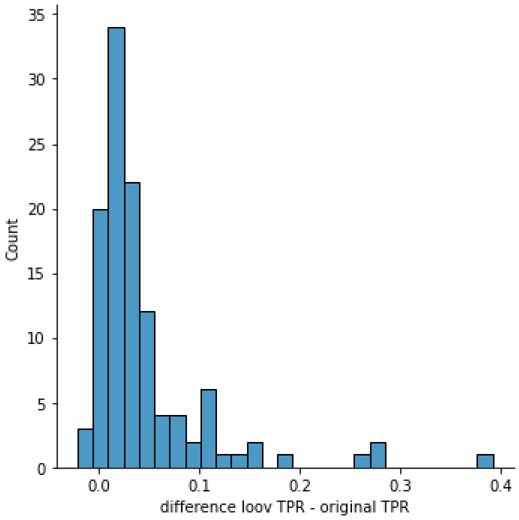

**Figure 3. Percentual TPR loss distribution after leave-one-out.**

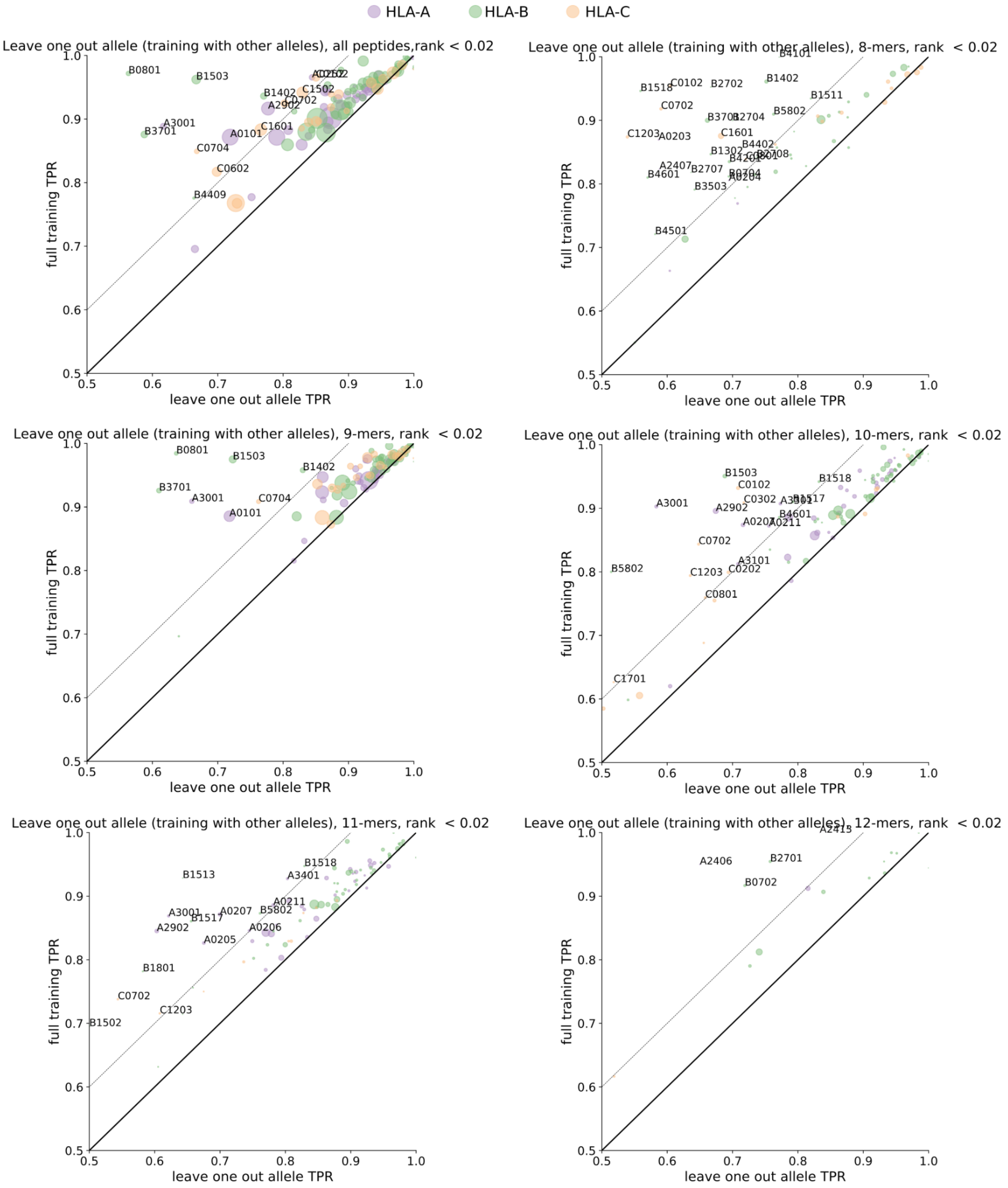

**Figure 4. HLA allele leave-one-out analysis.** The X-axis represents the leave-one-out allele TPR (by removing the allele peptides of the training set and re-training the model). The Y-axis represents the original TPR including all peptides in the training set. Thick continuous line overlaps with the diagonal. Dashed lines represent a %TPR loss greater or equal to 10%. HLA alleles with the annotated label are those with a %TPR loss greater than 10%.

#### 4. Neo expression adjusted model performance

We evaluated whether the expression adjusted model (see above Expression adjusted presentation likelihood) is able to improve the predictions from the expression naive model. To assess this, we leveraged the part of our training set that was obtained from ref.[2] Briefly, this is a publicly available dataset that contains experimentally identified peptides through immunopeptidomics across 95 HLA monoallelic cell lines engineered from B721.221. We therefore matched these observations with the RNA-seq expression of the source HLA-null cell line B721.221.

We observed that the model adjusted by the RNA expression has consistently higher TPR across all the evaluated ranking thresholds ($1e^{-06}$, $1e^{-05}$, $1e^{-04}$, $5e^{-04}$, $1e^{-03}$, $1e^{-02}$, $2e^{-02}$ and $1e^{-01}$, see Figure5 below). The average TPR across these thresholds was higher in the expression adjusted group (0.52) compared to the non-adjusted (0.45). Taken together, these observations confirm the added value of adjusting by the RNA expression of the source transcript. Unless otherwise specified, this model will be used to select the potential (neo)epitopes derived from tumor specific alterations in our neoepitope prioritization pipeline.

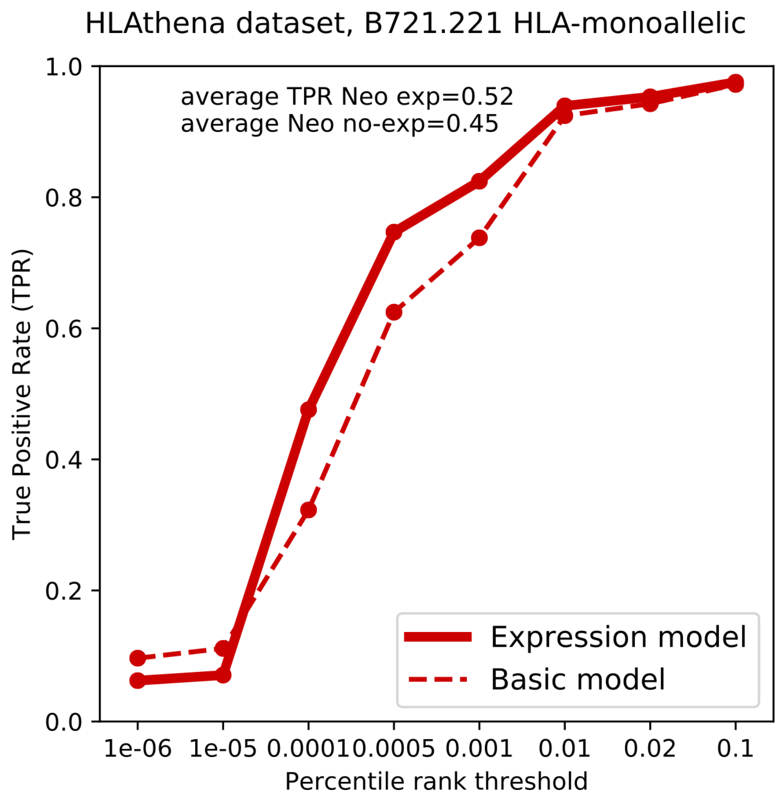

**Figure 5.** Performance comparison of expression adjusted versus non adjusted models. The x-axis represents the evaluated percentile ranking thresholds. The y-axis represents the TPR at the given thresholds. Basic model is the non-expression adjusted model.
